# Multiplexed Perturbation Enables Scalable Pooled Screens

**DOI:** 10.1101/2025.08.14.669942

**Authors:** Stefan Oberlin, Neil Tay, Albert Xue, Harold Pimentel, Michael T McManus

## Abstract

CRISPR-based genetic perturbation screens have revolutionized the ability to link genes to cellular phenotypes with unprecedented precision and scale. However, conventional pooled CRISPR screens require large cell numbers to achieve adequate sgRNA representation, posing technical and financial challenges. Here, we investigate the impact of co-delivery of multiple guide RNAs via high multiplicity of infection (MOI) in pooled CRISPR interference (CRISPRi) screens as a strategy to enhance screening efficiency while reducing cell numbers. We systematically evaluate screen performance across varying MOIs, assessing the effects of multiplexing on knockdown efficiency, sgRNA representation, and potential interference of multiple sgRNA phenotypes. Our data demonstrate that sgRNA multiplexing (MOI 2.5-10) can maintain screen performance while enabling significant reductions in cell number requirements. We further apply these optimized conditions to conduct a genome-wide CRISPR screen for regulators of the intracellular adhesion molecule ICAM-1, successfully identifying novel candidates using as few as half a million cells. This study provides a framework for adopting multiplexed sgRNA strategies to streamline CRISPR screening applications in resource-limited settings.

## Introduction

Genetic perturbation screens are a cornerstone technology to identify links between genes and cellular phenotypes^1,2^. CRISPR screens have drastically increased the accessibility and precision of these experiments and facilitated the discovery of countless novel biology^3,4^. Cas effector proteins are employed to knock out or modify the expression of genes or regulatory elements in a large population of cells. Changes in cell behavior are correlated with subsets of these genes linking genotype to phenotype. CRISPR screens rely on three essential components. First, cells need to express the Cas effector protein effectively. In many cases, stable cell lines are generated to ensure sufficient and homogenous expression of these enzymes, but transient expression has proven to a valid alternative. Secondly, single guide RNA (sgRNA) targeting a defined set of genes are expressed in these cells. sgRNA are usually delivered using lentiviral transduction to facilitate their integration into the host cell’s genome. This guarantees continued expression of sgRNA and allows for simple sequencing of the sgRNA populations directly from genomic DNA. Guide RNA are generally introduced so that a single copy is expressed from each cell. For lentiviral delivery, this is achieved by a low multiplicity of infection (MOI) of around 0.3-0.5, where in this case the MOI refers to the average number of sgRNA delivered and expressed per cell. This typically ensures that only one gene is perturbed per cell.

Thirdly, a very important parameter that affects screen performance is guide RNA representation or coverage. This metric represents the average level of replication per sgRNA in the screen. A higher average replication level enhances screen performance, with many labs aiming for a 200 to 500-fold coverage^5–7^. While CRISPR screening platforms are now widely available, aiming for these large replication levels poses many technical and financial challenges. To illustrate, around 50-100 million cells are required to fulfill these optimal conditions for a genome wide CRISPR screen. Maintaining these cell numbers for prolonged times can be problematic or even prohibitive in many cases such as when working in primary tissue or in vivo models. And with decreasing costs for sequencing, costs and labor required for cell growth and selection or isolation of a subpopulation of cells are gradually becoming the main obstacle to perform a CRISPR screen. This consequently heightens the incentives to develop more compact screening strategies. Several such strategies to compress CRISPR screens were developed and allow for the growth and selection of fewer cells. They aim at increasing the CRISPR efficiency by using the best performing sgRNA or by co-expressing multiple sgRNA against the same gene^8–10^. Furthermore, exceptionally homogenous sgRNA libraries and additional barcoding strategies were shown to improve screen performance with fewer cells^11–14^. An alternative strategy for screen compression is to perform it at higher MOI where multiple sgRNAs are expressed in each cell, or so to say under sgRNA multiplexed conditions. With the argument that multiple sgRNAs add noise or add to nonspecific toxic effects, this approach has been neglected. However, in specialized cases where most guide RNAs fail to induce an effect^15^, for example for CRISPR base editing screens^16^ or to study genetic interactions^17^, it may offer the same benefit.

Numbers of analyzed cells are even more so a limiting factor in high-content CRISPR screens, which are now available to directly link given phenotypes with induced perturbations at the single cell level^18^. For example, in single-cell CRISPR-seq (scCRISPR-seq) experiments, mRNA expression changes are directly linked to gene perturbations at scale^19,20^. Here, screen compression can be achieved by multiplexing of sgRNA expression cassettes using high MOI infections^21^. Random sgRNA combinations and their associated transcriptomes are then measured in single cells and subsequently deconvolved into gene expression changes associated to each perturbed gene. Similar success was achieved for a chemical screening platform using high-content imaging and scRNA-seq readouts^22^.

Given the success of high MOI infections for single cell approaches, we asked what is the effect of performing a classical pooled CRISPR screen under these multiplexed conditions? We therefore set out to systematically answer how multiplexing sgRNA delivery through high MOI impacts pooled CRISPR interference (CRISPRi) screen performance. We focused on comparing performance across varying MOIs and establish guidelines for the maximal compression potential by reducing cell numbers across conditions. Anticipating loss of CRISPRi activity upon high degrees of multiplexing and potential performance decrease through interference of multiple sgRNA phenotypes, we assayed multiplexing performance at a single cell level for a select set of target genes and subsequently benchmarked performance in a pooled cell essentiality and drug tolerance screen. We ultimately made use of these optimized conditions to perform a compressed, pooled, genome-wide CRISPR screen identifying novel regulators of the intracellular adhesion molecule ICAM-1 by sorting as little as half a million cells.

## Results

### High MOI infection allows for multiple gene repression

To evaluate the feasibility of CRISPR screening compression via high multiplicity of infection (MOI), we first sought to ensure that gene perturbation occurs efficiently when multiple sgRNAs are introduced into cells (**Fig. 1A**). To achieve this, we began by optimizing lentiviral infection conditions and quantifying infection efficiency, which is critical for accurate guide RNA (gRNA) delivery via lentiviral transduction. The gold standard for measuring DNA copy numbers introduced per cell is digital PCR (dPCR), a highly sensitive and quantitative technique^23^. To determine the optimal infection conditions, we used the widely employed spinfection protocol in K-562 cells, a chronic myeloid leukemia cell line. Spinfection resulted in a 2- to 5-fold increase in infection efficiency compared to conditions without centrifugation or polybrene supplementation (**Supplementary Fig. 1A-B**). Interestingly, the degree of improvement varied depending on the viral dose used, suggesting that infection efficiency does not scale linearly with the volume of virus added.

**Figure 1:**
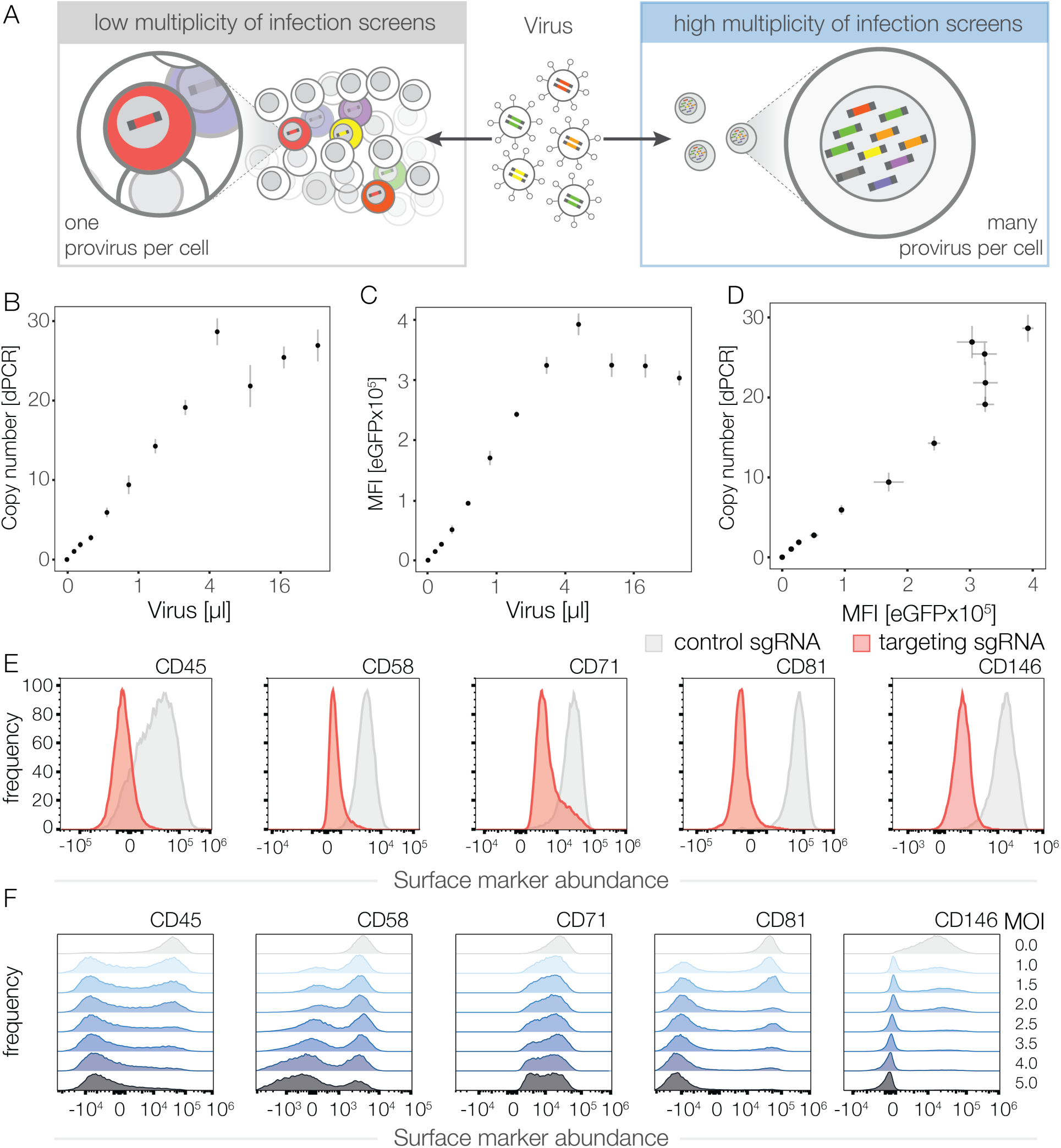
Lentiviral high MOI CRISPR guide RNA delivery and simple copy number quantification. **A)** Illustration of low and high multiplicity of infection (MOI) lentiviral sgRNA delivery in CRISPR applications. **B)** Quantification of lentiviral insertions relative to viral concentration using dPCR from genomic DNA. **C)** eGFP expression (MFI, mean fluorescent intensity) from Puro-T2A-eGFP marker cassette used in sgRNA expression vectors correlates with quantity of transduced lentivirus. **D)** Correlation between fluorescent marker expression and dPCR-inferred copy numbers of lentiviral insertions. **E)** CRISPRi knockdown levels in K-562 cells with a single sgRNAs targeting five surface markers individually, measured by antibody staining and flow cytometry. **F)** Knockdown efficiency using pooled sgRNAs against five surface markers at increasing MOI. In B-D dots represent the mean of three replicates and bars the standard deviation from the mean.

To investigate these results further, we infected cells with a range of viral concentrations and quantified the resulting copy number per cell using dPCR. The data revealed a steady increase in infection levels with higher viral inputs, but saturation occurred at approximately 30 copies per cell (**Fig. 1B**). This saturation effect raised questions about potential biological and physical limitations of lentiviral transduction. It is plausible that cellular entry mechanisms or intracellular processing capacity may impose constraints on the number of viral particles that can be successfully integrated. Notably, we observed that different viral preparations resulted in varying MOI saturation points, with higher-quality viral preps achieving greater copy numbers.

However, dPCR is a relatively time-consuming procedure that requires specialized instrumentation, making it inaccessible to many laboratories. To address this limitation, we explored alternative strategies that could provide reliable estimates of lentiviral infection levels in a more practical and accessible manner. Lentiviral vectors used for sgRNA delivery, including ours used in this study, typically include an antibiotic resistance gene for selection purposes and, in some cases, a fluorescent marker such as enhanced green fluorescent protein (eGFP) to facilitate cell tracking. We therefore measured mean fluorescence intensity (MFI) via flow cytometry (**Fig. 1C**). The MFI values closely mirrored the trends observed with dPCR measurements, with a similar saturation pattern emerging at higher viral doses (**Fig. 1D**). This linear relationship between MFI and viral copy number suggests that MFI can serve as a practical and reliable surrogate for estimating infection efficiency in experimental settings, offering a streamlined and cost-effective alternative to dPCR. Furthermore, control cells infected at a low MOI (∼0.3), where most infected cells harbor a single sgRNA insertion, can serve as a simple reference standard for linear extrapolation of fluorescence signals to estimate MOI across different conditions.

Given the potential for large-scale genome rearrangements and unintended off-target effects associated with high-order Cas9 nuclease-based perturbations, we opted to utilize a CRISPR interference (CRISPRi) approach. Here we employed the catalytically inactive Cas9 (dCas9) fused to the Zim3 KRAB repressive domain to achieve targeted gene silencing^24^. To evaluate the performance of this system, we infected K-562 CRISPRi cells with sgRNAs targeting five distinct cell surface markers individually. Flow cytometry analysis confirmed nearly complete knockdown of the target proteins, validating the effectiveness of single sgRNA perturbations (**Fig. 1E**, **Supplementary Fig. 1C**).

Aiming to test whether expression of multiple sgRNA yields efficient simultaneous gene repression, we then proceeded to pool the five sgRNAs and performed multiplexed infections at increasing MOIs. To quantify knockdown efficiency accurately, we calculated the probability of targeting multiple surface markers in individual cells based on MOI (**Supplementary Fig. 1D**). It is important to note that actual copy numbers per cell follow a Poisson distribution around the averaged MOI. Our calculation also takes into consideration that at higher MOI multiple copies of the same sgRNA can be present. As expected, the extent of surface marker repression scaled with higher MOIs, with all five markers exhibiting significant knockdown at elevated MOI levels (**Fig. 1F**). The distribution of the number of repressed genes in each cell also scales with increasing MOI and all five genes are simultaneously repressed in about 42% of cells at the highest MOI of five where 56% of cells are expected to carry five or more sgRNA copies (**Supplementary Fig. 1D-E**). This suggest that around three quarter of cells with five or more sgRNA copies repress all five surface markers. The same trend was observed comparing the fraction of expected versus observed quintuple knockdowns across varying MOIs (**Supplementary Fig. 1F**). Collectively, our results suggest that high MOI infections are easily quantifiable by flow cytometry when using sgRNA vectors with fluorescent protein expression and allow for the simultaneous repression of at least up to five genes. To ensure the robustness and universality of our approach, we replicated these experiments in HEK-293T cells and observed highly consistent results, confirming that our optimized infection strategy is applicable across different cell lines (**Supplementary Fig. 2**).

### Developing a Compact sgRNA Library for Benchmarking High MOI Pooled CRISPRi Screens

Our initial data suggested that multiplexed CRISPRi with high MOI sgRNA delivery could significantly enhance the efficiency and scalability of pooled CRISPR screens. To further investigate the potential of this method, we designed a compact screening protocol that closely emulates commonly used genome-wide CRISPR libraries, enabling the assessment of screening performance across various experimental conditions. For this purpose, we designed a compact 2000 sgRNA library targeting epigenetic regulators (**Supplementary Fig. 3A-C**). The library construction involved cloning the sgRNAs into an expression vector that contained a semi-random barcode, allowing for unique identification of each genomic sgRNA insertion. Each gene in the library is targeted by three independent sgRNAs to enhance targeting robustness, and 5% of non-targeting sgRNAs, serving as internal controls to monitor background effects and non-specific responses.

Moreover, the fraction of essential genes closely mirrors the one observed in genome-wide libraries. It therefore offers a streamlined yet comprehensive approach for a functional genomic study and for assessing the impact of CRISPRi perturbations in pooled screens. Furthermore, epigenetic regulators are largely expressed in K-562 cells (**Supplementary Fig. 3D-E**) and are known to influence drug resistance mechanisms^26^, making them particularly relevant for our experimental objectives.

### Evaluating the Influence of sgRNA Infection Levels on Pooled CRISPRi Screen Performance

Using this compact sgRNA library, we aimed to evaluate the impact of sgRNA multiplexing through high multiplicity of infection (MOI) transductions in a pooled CRISPR screen. We measured both cell essentiality as well as drug tolerance against the tyrosine kinase inhibitor Imatinib (**Fig. 2A**). The CRISPR screen was conducted under three experimental conditions designed to assess the effects of MOI and cell number on screen performance (**Fig. 2B and Supplementary Table 1**). To ensure sufficient representation of the library, cell populations were maintained at a minimum representation of 250× as a baseline. In the constant cell numbers condition, cells were maintained at consistent cell numbers across all MOI conditions, leading to an increase in the number of sgRNAs proportional to the MOI. This condition tests the effect of increasing the MOI under ample cell representation. In the constant sgRNA numbers condition, the number of cells per passage was adjusted to achieve a constant total number of sgRNAs across MOI conditions; for example, five times fewer cells were passaged at MOI 5 compared to the single insertion reference. Here, we aimed to identify how much increasing the MOI can compensate the reduction of cells. Lastly, in the low MOI cell number control condition, cells were infected at a fixed low MOI (0.3) but passaged at cell numbers equivalent to the constant guide RNA numbers condition.

**Figure 2:**
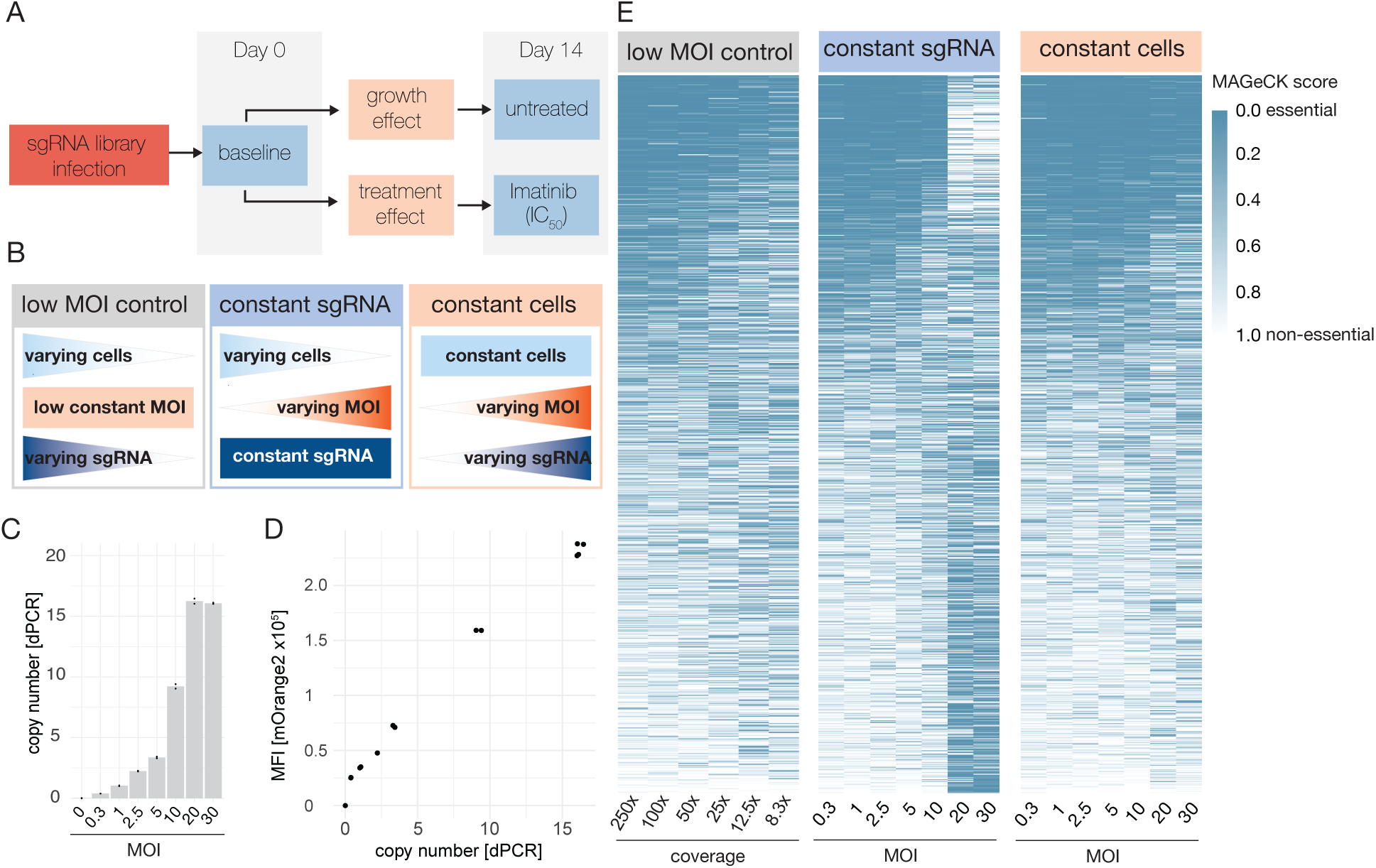
MOI-dependent CRISPRi efficacy in genetic screens for essential gene and drug tolerance. **A)** Schematic workflow of CRISPRi screens for cell essentiality and drug tolerance assessment. **B)** Experimental conditions for evaluating lentiviral MOI effects on CRISPRi screen performance. **C)** A compact sgRNA library targeting epigenetic regulator genes was transduced into K-562 CRISPRi cells with varying MOI and lentiviral insertions are quantified by dPCR from genomic DNA. **D)** Correlation between mOrange2 fluorescence intensity, expressed from the sgRNA expression vector, and lentiviral copy numbers determined by dPCR. **E)** Gene essentiality scores calculated via MAGeCK RRA algorithm across the three experimental conditions described in **(B)**.

K-562 cells were transduced in duplicate under varying infection levels, ranging from MOI 0.3 to 30 (**Fig. 2C**). After puromycin selection to enrich for transduced cells, we validated sgRNA expression cassette copy numbers using digital PCR (dPCR). Notably, cells infected at the two highest MOIs exhibited copy number plateaued slightly below their expected values (**Fig. 2C**). Additionally, fluorescence signals of the mOrange2 reporter correlated linearly with the copy numbers measured via dPCR, further confirming successful sgRNA multiplexing (**Fig. 2D**).

Because of the uniquely barcoded sgRNAs, we couldtrack individual infection events by sequencing. This confirmed the anticipated diversity of barcode-to-sgRNA combinations across all conditions at the start of the screen (**Supplementary Fig. 3F**). Cells were cultured for 14 days under untreated conditions or treated with imatinib (IC50: 230 nM), a tyrosine kinase inhibitor targeting the constitutively active BCR-ABL fusion protein caused by the Philadelphia chromosome^25^. This K-562 model system has been widely used in CRISPR-based screens to investigate mechanisms of drug resistance^26–28^. By benchmarking our experimental setup across the three conditions, we aimed to evaluate the performance of high MOI screens and determine the feasibility of compressing screens by reducing cell numbers. To estimate guide RNA level effect sizes, we employed DESeq2^29^ and to estimate gene level effect sizes, we employed MAGeCK robust rank aggregation^30^. MAGeCK scores at the gene level indicate that cell essentiality estimates largely remain stable across different conditions, with some specific exemptions (**Fig. 2E**). Together, these data illustrate a robust method for testing the scalability and efficiency of highly multiplexed pooled CRISPRi screens.

We evaluated the ability of screens at different MOIs to identify cell-essential and cell-nonessential genes using prior high-quality classifications^31^ as a benchmark (**Fig. 3A**). The sgRNA level effect sizes are then ranked and the area under the curve (AUC) for cell essential genes serves as a first assessment of screen performance, here for example under standard low MOI conditions (**Fig. 3B**). We evaluated the classification performance at the gene level by calculating the receiver operating characteristic area under the curve (ROC-AUC), using essential genes as positive controls and non-essential genes as negative controls. These estimates were then used to compare different conditions at the gene level as exemplified by lower performances when reducing the passaged cell numbers in the low MOI control (**Fig. 3C**). Similarly, in the constant cell number condition, we could identify the impact of MOI alone on screen performance (**Fig. 3D-E)**, suggesting that the MOI ranges of 2.5–10 are ideal for high-throughput CRISPRi screens. These finding were further corroborated by analyzing sgRNA-level accuracies and inter-replicate correlations (**Supplementary Fig. 4 A-B**). We also examine how guide RNA effect sizes change with increasing MOI. On average, effect sizes become more positive with higher MOI when considering all genes or cell essential genes only, but not for non-essential genes (**Supplementary Table 2**). These results indicate that increasing MOI moderates essential gene effect estimates, which are inferred with a bias with low initial counts.

**Figure 3:**
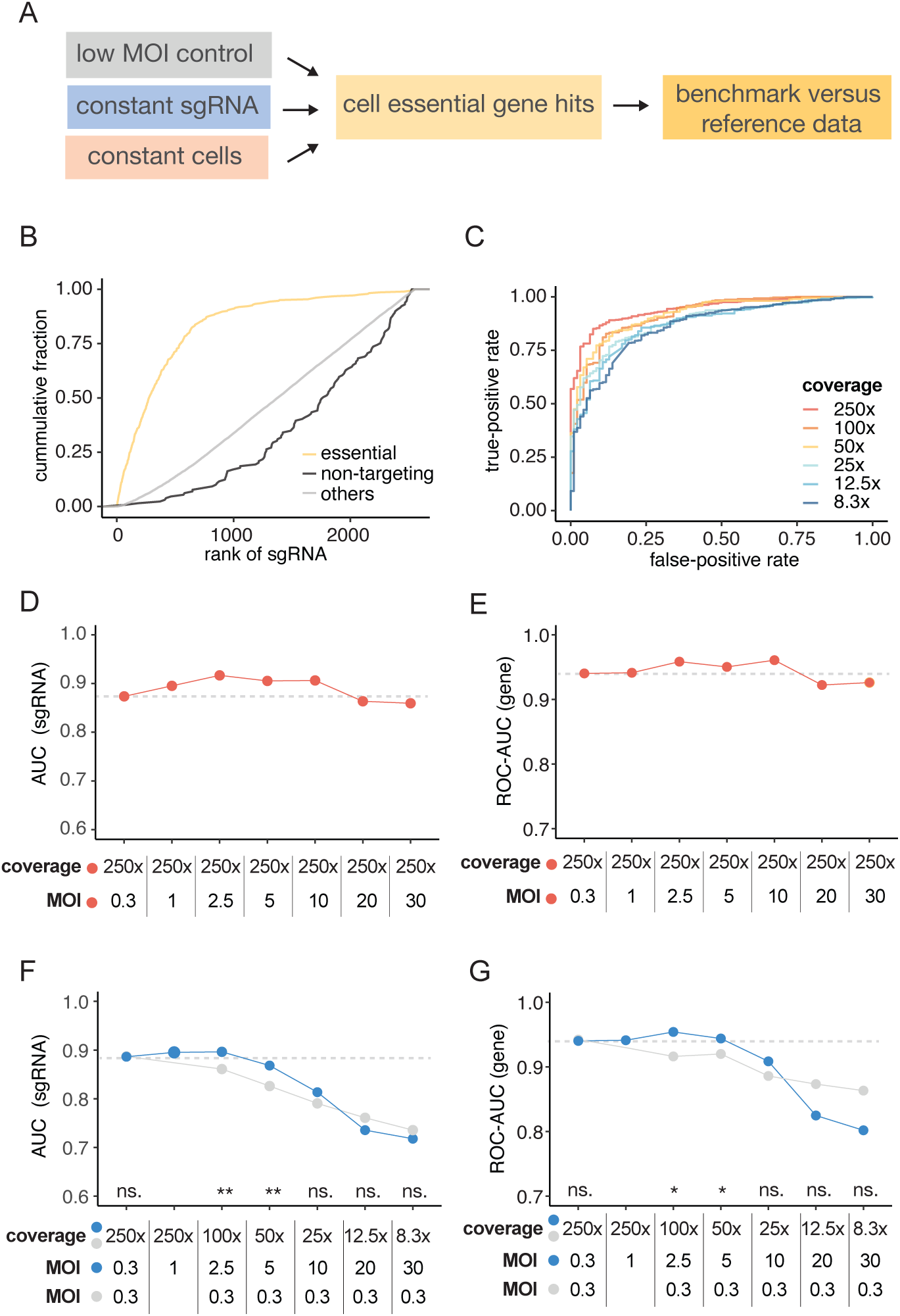
Guide RNA libraries at moderately high MOI improve CRISPRi screen performance. **A)** Workflow for essential gene identification under variable CRISPRi screen conditions and subsequent performance evaluation. **B)** Ranked cumulative distribution comparing sgRNA targeting essential genes (from Horlbeck et al. 2016), non-targeting controls and other gene targets in the MOI 0.3, 250x cell coverage condition. The cumulative distribution of cell essential gene targeting sgRNA is used to calculate the area under the curve (AUC). **C)** AUC for the receiver operating characteristic (ROC) analysis of MAGEcK gene scores for cell essential gene identification in the constant cells condition. **D)** Comparison of ranked sgRNA cumulative distribution AUC and **(E)** MAGEcK gene score ROC-AUC for essential gene identification across MOIs in the constant cells condition (red). **F)** Comparison of ranked sgRNA cumulative distribution AUC and **(G)** MAGEcK gene score ROC-AUC for essential gene identification across MOIs in both constant sgRNA (blue) and low MOI control (grey) conditions. DeLong test. \**P* < 0.05, \*\**P* < 0.01, ns. non-significant.

### High MOI can partially compensate for reduced cell numbers in CRISPRi screens

Having established that CRISPRi screen performance for identifying cell-essential genes is optimal at medium-high guide RNA infection levels, we next sought to determine to what extent higher MOIs can compensate for performance loss when reducing cell numbers. To evaluate this, we again performed AUC analysis on sgRNAs ranked by their level of depletion in the screen. We primarily compared the constant sgRNA and the low MOI control condition to control for the passaged cell numbers. Confirming our prior findings, AUC analysis at sgRNA-level remains constants at low to medium MOIs of 0.3 to 5 in the constant sgRNA condition (**Fig. 3F**). By contrast in the low MOI control, AUC analysis reveals a steady decline in performance for reduced cell numbers. This drop begins at a 2.5-fold reduction in cell number (100x cell coverage) and continues to decrease with greater reductions, culminating in a substantial drop at a 30-fold reduction in cell numbers (**Fig. 3F**). This result aligns with previous observations that lower cell representation negatively impacts CRISPRi screen performance^5,32^. Guide RNA-level accuracies and inter-replicate correlations analysis across all conditions supports these findings (**Supplementary Fig. 4C-D**), further highlighting that increasing MOI can partially compensate for reduced cell numbers, provided the MOI remains within the range of 2.5 to 5.

To assess screen performance at the gene level, guide RNA effects were again aggregated into gene-specific phenotypic scores. ROC-AUC analysis of these gene-specific scores for cell-essential genes mirrored the sgRNA-level results. In the constant cells condition, ROC-AUC remains relatively unchanged from low to medium MOIs (**Fig. 3G**). Reducing cell numbers in the low MOI control led to a gradual decrease in ROC-AUC. Interestingly, performance at high MOI of 20 to 30 drops precipitously when simultaneously reducing cell numbers. This outcome is likely due the very low cell numbers evaluated combined with high guide RNA collision rates or reduced gene knock-down at high MOI.

It is expected that less abundant sgRNA in the original plasmid library perform poorer than more abundant sgRNA. These less abundant sgRNA are more likely to drop out of the screen in case of a bottlenecking event independent of their actual effect size. We therefore stratified the sgRNA into quartiles based on their abundance levels in the plasmid library (**Supplementary Fig. 5A**). Note, these abundances are independent of sgRNA effect size and merely reflect the clonability of each sgRNA sequence. Interestingly, screen performance improvements in higher MOI conditions are most pronounced in lower abundant sgRNA (quartile1 and to a lesser degree quartile 2) and are almost absent in highly abundant sgRNA (quartile 3 and 4, **Supplementary Fig. 5B**). This suggests, unsurprisingly, that the performance increase is likely due to the decreased bottlenecking events.

Our results challenge the prevailing notion that CRISPR screens must be initiated at low MOI levels (e.g., 0.3–0.5). Instead, our findings suggest that performing CRISPRi screens with medium MOIs and large coverage is optimal for identifying or validating cell-essential genes. Furthermore, comparable performance can be achieved when screens are conducted with cell numbers reduced by 2.5- to 5-fold, provided that sgRNA expression is multiplexed to the same extent. Specifically, screens performed with a 50× guide RNA representation achieve results comparable to those widely recommended under traditional settings (e.g., 250× representation at MOI 0.3).

### Evaluating the Impact of MOI on Identifying Drug Tolerance Genes in CRISPRi Screens

We next evaluated the effect of high MOI guide RNA multiplexing in a drug resistance screen (**Fig. 4A**). Specifically, we assessed the ability to identify gene knockdowns that render K-562 cells resistant to BCR-ABL1 tyrosine kinase inhibition. Quantitatively evaluating high MOI multiplexing of a drug resistance CRISPR screen should be inherently more challenging than evaluating performance for identifying cell-essential genes. First, there are limited reliable ground truth data for comparison. Second, the percentage of true positive hits is significantly lower, making the screen more susceptible to noise. We thus reasoned that a drug resistance screen should set a higher benchmark.

**Figure 4:**
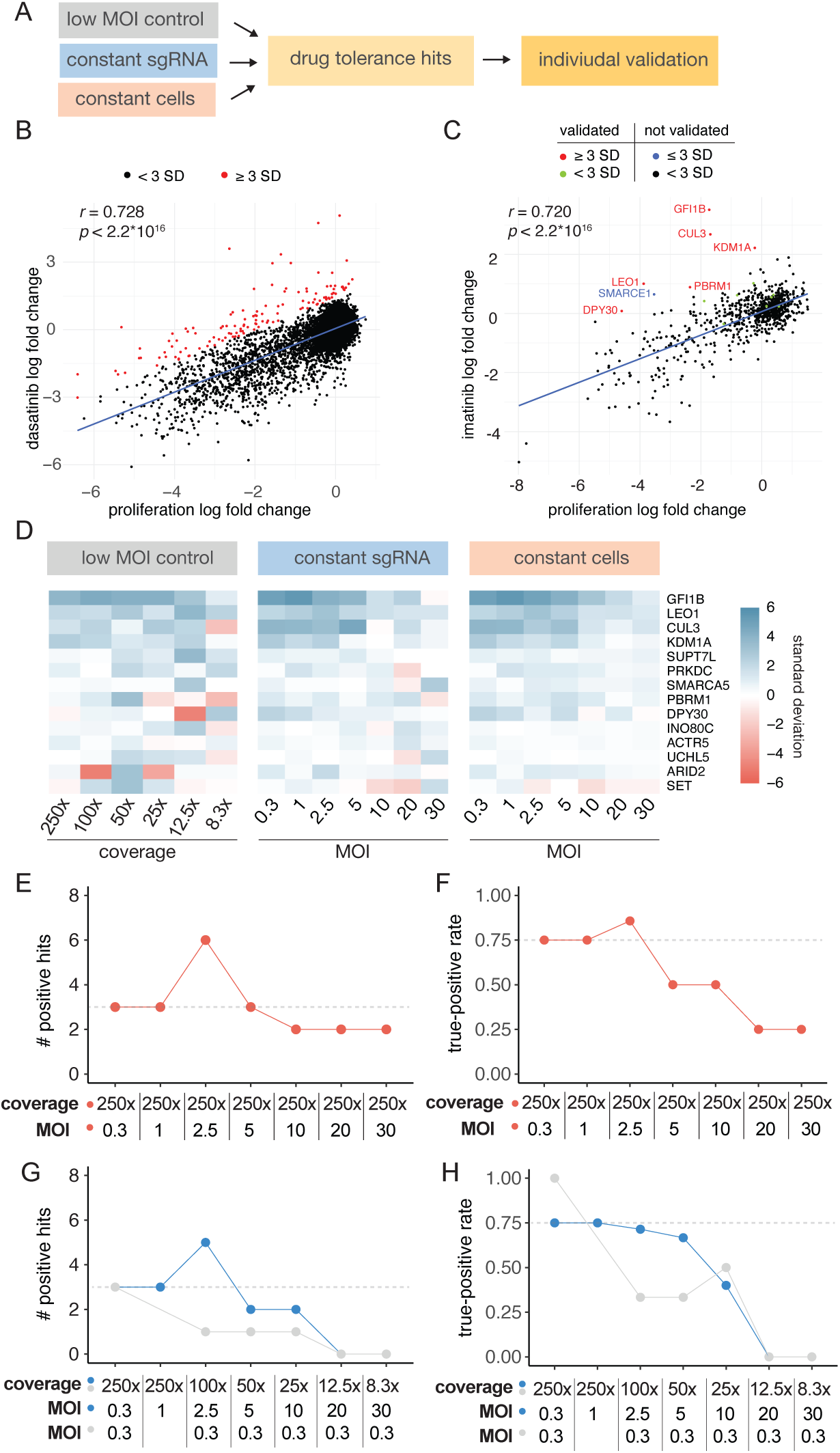
Impact of sgRNA multiplexing on CRISPRi drug tolerance screen. **A)** Workflow to identify drug tolerance genes under varied CRISPRi screen conditions and individual validation of drug tolerance hits to benchmark screen performance. **B)** Correlation of gene-level guide RNA proliferation and drug treatment effects from high quality reference tyrosine kinase inhibitor screen (Heo et al. 2024). MAGeCK cell proliferation and drug effect fold changes are plotted against each other highlighting the linear relationship between these two effects. Positively selected genes with residual >3 standard deviation are classified to have a drug tolerance effect (red points). **C)** Tyrosine kinase inhibitor screen performed with an epigenetic regulator library in the constant cell condition with an MOI 5. Individually validated genes with residual ≥3 standard deviations (red points) and <3 standard deviations (green points) in this screening condition, non-validated genes with residual ≥3 standard deviations (blue points) and <3 standard deviations (black points) in this screening conditions. **D)** Drug tolerance effects of all individually validated genes included in the epigenetic regulator guide RNA library in all three experimental conditions across varying MOI and cell coverage. **E)** Number of true positive drug tolerance hits in in the constant cells condition. **F)** True-positive rate of drug tolerance hits identification in in the constant cells condition. **G)** Number of true positive drug tolerance hits in in the constant sgRNA and low MOI control conditions. **H)** True-positive rate of drug tolerance hits identification in in the constant sgRNA and low MOI control conditions.

To establish a benchmark, we used a recent high-quality CRISPRi screen for tyrosine kinase inhibitor resistance performed in K-562 cells under dasatinib treatment^11^. Genes conferring resistance were identified by comparing MAGeCK scores between untreated and treated conditions against baseline measurements at the start of the screen. This comparison revealed a linear correlation between cell-essential gene knockdown effects and cell proliferation, with deviations from the correlation representing treatment-specific resistance phenotypes. A linear model was fit to the data, and genes deviating by more than three standard deviations from the model were classified as hits. This approach identified 130 high-confidence resistance genes from the genome-wide CRISPRi screen (**Fig. 4B**).

We then applied the same selection criteria to our dataset, restricted to epigenetic regulators (**Fig. 4C-D**). All of these hits were individually validated to confirm their effect on imatinib tolerance (**Supplementary Fig. 6**). First, we examined the overlap of resistance hits across MOI conditions in the constant cell number condition (**Fig. 4E-F**). As expected, the overlap with individually validated hits was highest at low to mid MOIs (0.3 to 5), with three resistance hits and true positive rates mostly exceeding 0.75. Interestingly, performance at MOI 2.5 was notably superior, identifying six resistance genes that overlapped completely with the validated hits.

Next, we compared screen performance in the constant guide RNA condition with the low MOI cell number control (**Fig. 4G-H**). Performance was comparable between standard low MOI conditions and 2.5- to 5-fold cell number reductions, but only when MOIs were increased accordingly. Conversely, reducing cell numbers by 2.5- to 5-fold without increasing MOI significantly impaired screen performance. This condition yielded low true-positive rates and failed to identify most positive hits.

Taken together, these results demonstrate that guide RNA multiplexing (2.5- to 5-fold) in a CRISPRi screen not only enhances the identification of cell-essential genes but also improves the detection of drug resistance genes. These findings highlight the potential of multiplexed guide RNA expression to optimize the performance of CRISPR screens, even under challenging conditions such as drug resistance profiling.

### Genome-wide CRISPRi Screen to Identify Regulators of CD54 (ICAM-1) Expression

To further explore the potential of guide RNA multiplexing in pooled CRISPRi screens, we performed a genome-wide screen to identify regulators of CD54 (ICAM-1) levels in K-562 CRISPRi cells (**Fig. 5A**). A compact barcoded dual-sgRNA library where each gene is targeted by a single vector expressing two sgRNA was used^8^. Cells were infected with the guide RNA library at two different MOIs (0.3 and 5), followed by puromycin selection to ensure successful transduction. To verify infection efficiency, we measured the mean fluorescence intensity (MFI) of mBFP2 in cells infected at both MOI 0.3 and 5 (**Supplementary Fig. 7A**). After selection, cells were stained with an anti-CD54 antibody and sorted into three distinct populations based on CD54 expression levels – low and high (each representing 10% of the population) and mid (20%). Analytical flow cytometry of the sorted cell populations further confirmed the accurate separation of cells based on CD54 expression levels (**Fig. 5B**). The experiment was conducted in duplicates at three different cell coverage levels (25X, 125X, and 250X cell coverage) to rigorously evaluate the impact of cell number representation on screen performance.

**Figure 5:**
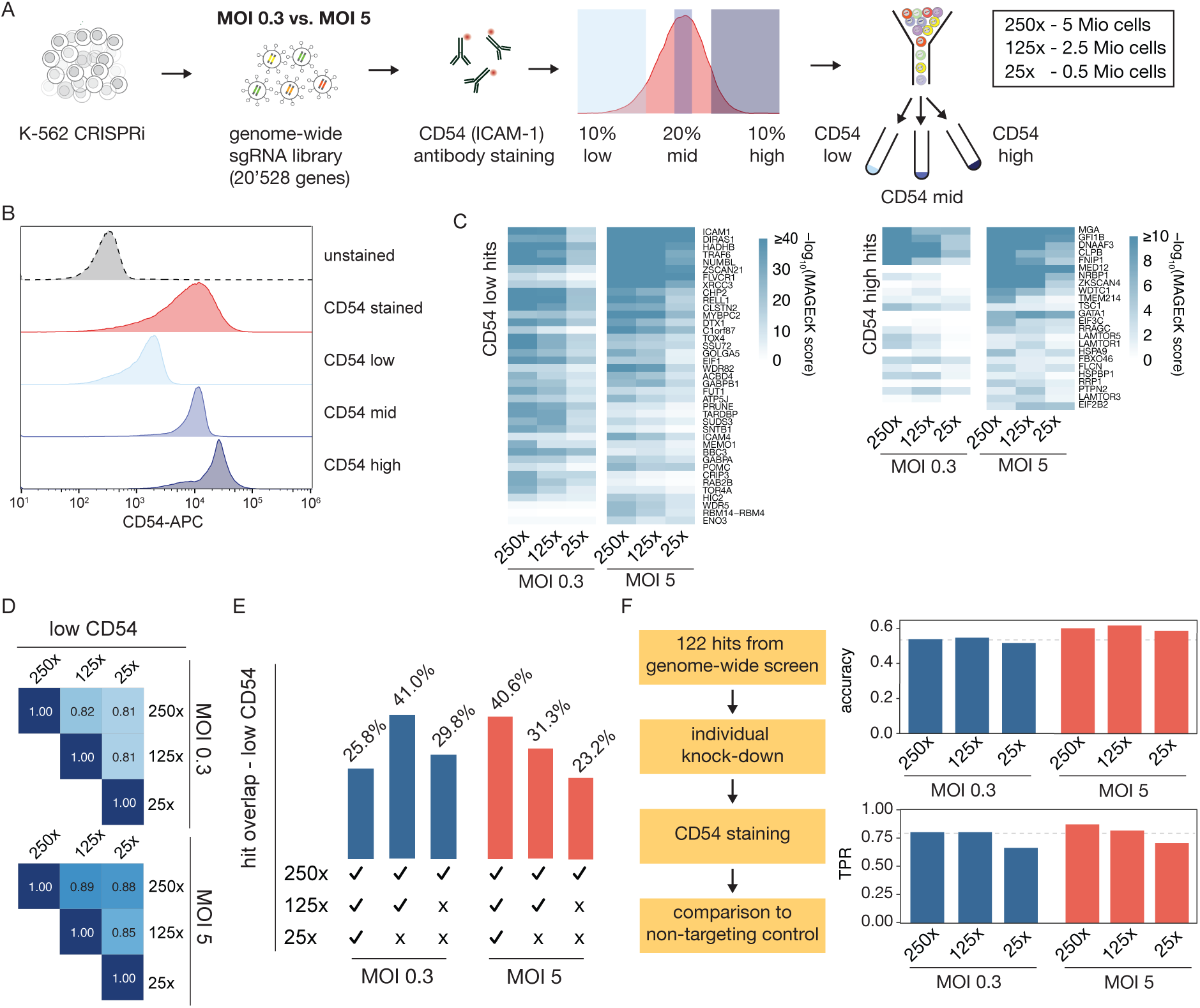
Ultra-compressed FACS sorting CRISPR screen with guide RNA multiplexing. **A)** Schematic of genome-wide CRISPRi screen identifying CD54 (ICAM-1) regulators. K-562 CRISPRi cells were infected with a guide RNA library at MOI 0.3 and 5, then sorted into three bins (low/high 10%, mid 20%) based on CD54 antibody staining. The screen was conducted at three different library representation levels. **B)** Analytic flow cytometry of sorted cell populations. **C)** CRISPRi screen hits showing positive regulators (CD54 low bin) and negative regulators (CD54 high bin) of CD54. MAGeCK scores across all experimental conditions are displayed for individually validated hits. **D)** Pearson correlation of samples with varying cell coverage, based on log_2_ fold changes of normalized sgRNA abundances between low and mid CD54 populations. **E)** Percentage of shared hits between samples with varying cell coverage within the MOI 0.3 and MOI 5 condition. **F)** Hits selected across all conditions were individually validated to calculate true positive rate (TPR) and accuracy to identify hits.

Screen enrichment analyses were performed to identify sgRNAs enriched in cells with altered CD54 expression. Using our barcoded sgRNA library, we applied MAGeCK to identify gene-level scores in CD54 low samples (positive ICAM-1 regulators) and CD54 high samples (negative ICAM-1 regulators) (**Fig. 5C**). Genes with a false discovery rate (FDR) below 0.01 were considered significant hits. Notably, the analysis identified several strong regulators of CD54 expression, such as the NF-κB activator TRAF6 as well its interaction partner AMBRA1 and NUMBL, known to regulate ubiquitination of TRAF6^33,34^, with distinct enrichment patterns observed across different MOI and cell representation conditions. To evaluate the consistency of sgRNA enrichment across experimental settings, we performed comparative analyses contrasting low versus mid CD54 expression samples. A first glance at screen performance is granted by ICAM-1 targeting sgRNA (**Fig. 5C**, **Supplementary Fig. 7B**) that are, expectedly, significant hits in all conditions. Pearson correlation analysis based on gene-level MAGeCK fold changes, restricted to genes identified as significant hits in any condition, confirmed greater consistency between high MOI samples across all cell coverages (**Fig. 5D, Supplementary Fig. 7C**). Euclidean distance calculations across all sgRNA fold changes further corroborate these results (**Supplementary Fig. 7D**).

We next classified screen hit overlaps between MOI conditions, dependent on cell coverage and identified higher overlaps between high and low coverage samples in high MOI compared to low MOI conditions (**Fig. 5E, Supplementary Fig. 7E**). To gain deeper insights into hit enrichment patterns, we stratified genes identified in the 250x condition into quartiles based on their MAGeCK enrichment scores (**Supplementary Fig. 7F-G**). Quartile 1 represented the most highly enriched hits, while quartile 4 included the least enriched hits. The FDR distribution remains low in quartile 1 of all conditions, indicating that the strongest hits are identified independent of MOI and cell coverage. FDR distributions in quartile 2 and more pronounced in quartiles 3 and 4 are consistently lower at MOI 5 at lower cell coverage. Taken together, these data suggest that screens performed at moderate MOI outperform standard low MOI screen with low cell numbers in particularly at identifying moderately enriched screen hits.

To validate the screen data, we performed secondary arrayed validation experiments on selected hits making sure that screen hits from all conditions are equally represented (**Supplementary Fig. 7H**). By comparing validated hits to their original screen classifications, we calculated screen accuracy across conditions, revealing that MOI 5 with high cell numbers achieved the highest accuracy and true positive rate (**Fig. 5F**). Accuracy remained high with lower cell numbers at MOI 5. In contrast accuracy at MOI 0.3 was lower and is more severely impacted by reducing cell numbers in the screen. Collectively, these data suggest that performing a genome-wide CRISPR at 25x cell coverage, or around half a million sorted cells, yields a screen performance comparable to current standard screening modalities.

To gain deeper insight into the functional roles of genes identified in our genome-wide CRISPRi screen, we performed a network-based analysis of known interactions between genes that positively and negatively regulate ICAM-1 (CD54) expression. The identified genes were categorized based on their regulatory effects, with positively regulating genes shown on the left and negatively regulating genes on the right (**Supplementary Fig. 8**). Network analysis revealed distinct clusters of functionally related genes within both positive and negative regulators of ICAM-1. Among the positively regulating genes, key pathways associated with immune signaling, transcriptional regulation, and Histone H4-K16 acetylation were enriched. The network highlights the central role of key transcription factors and signaling proteins in enhancing ICAM-1 expression. Conversely, negatively regulating genes formed distinct functional clusters associated with translation initiation, mTOR signaling and transcription factors regulating hematopoietic stem cells. Notably, two phosphatases PTPN1 and PTPN2 are strong negative regulators of ICAM-1. Both were previously identified to dampen NK cell killing in their target cells by inhibiting IFNγ signaling^35^. Our results suggest that increased immune synapse interaction through upregulation of ICAM-1 could cause this enhanced immune response upon PTPN1 or PTPN2 downregulation.

Our data provide a comprehensive overview of the molecular mechanisms governing ICAM-1 expression and identify key regulatory networks that positively and negatively influence its levels. The identification of genes associated with distinct functional pathways suggests potential targets for modulating ICAM-1 in therapeutic contexts, such as immune modulation and cancer therapy. Further studies leveraging these data could provide new avenues for targeted intervention in diseases where ICAM-1 plays a critical role.

## Discussion

In this study, we systematically evaluated the impact of guide RNA multiplexing via high multiplicity of infection (MOI) in pooled CRISPR interference (CRISPRi) screens to improve efficiency while reducing the number of required cells. Our data demonstrate that moderate MOI levels (2.5-10) not only maintain screen performance but can enhance the identification of both cell-essential and drug resistance genes when paired with appropriate cell numbers. These insights provide a framework for optimizing pooled CRISPR screen design, enabling resource-efficient approaches without compromising sensitivity and specificity.

One of the key data of this study is the nonlinear relationship between viral load and infection efficiency, where infection levels plateau at approximately 30 viral copies per cell. This saturation effect highlights the importance of optimizing MOI to avoid excessive viral integration, which can introduce confounding factors such as cellular toxicity and reduced knockdown efficiency. Our data further suggest that fluorescence intensity (MFI) can serve as a reliable proxy for assessing infection efficiency, offering a simpler alternative to more labor-intensive methods such as digital PCR.

We also demonstrated that high MOI CRISPRi screens with multiplexed sgRNAs can significantly reduce the negative impact of cell number reductions. Screens conducted at MOI 2.5-5 outperformed many standard low MOI (0.3) conditions, suggesting that higher MOI can partially compensate for lower cell numbers by increasing the effective coverage of sgRNAs. However, our results indicate that excessive sgRNA multiplexing (MOI >10) can lead to diminished performance, likely due to increased collision effects and off-target interactions that complicate data interpretation.

Our data highlight that guide RNA multiplexing can be effectively integrated with other screening compression strategies, such as the use of dual guide RNA libraries. These compression strategies can greatly contribute to cost reductions while maintaining robust screen performance. This includes reductions in cell culture reagents and selection reagents used for drug treatments or cell sorting, with applicability to any labor-intensive and costly methods for enriching specific phenotypes. Thanks to its simplicity and accessibility, our approach can be readily adopted by any laboratory performing CRISPR screens.

The potential to extend our data to in vivo applications presents exciting opportunities for translational research. High MOI multiplexing strategies could facilitate efficient perturbation of target genes in complex biological systems, such as animal models and patient-derived xenografts. However, additional considerations such as immune responses to viral delivery, off-target effects, and tissue-specific variability must be addressed to ensure the feasibility and reproducibility of in vivo applications. Further optimization of delivery methods, such as lipid nanoparticles or non-integrating viral vectors, may expand the applicability of these approaches in preclinical and clinical settings.

Despite the promising data of this study, there are limitations to consider. First, the observed improvements in screen performance may be context-dependent and require validation across diverse cell types and experimental conditions. High MOI multiplexing can also introduce challenges such as increased rates of off-target effects and cellular toxicity due to excessive viral integration. Researchers may find an asset for high MOI multiplexing when studying cellular phenotypes in primary cells. While our study focused on model cell lines, the applicability of high MOI multiplexing in primary cells and in vivo settings where factors such as transduction efficiency and cell heterogeneity can impact outcomes. We implore investigators to conduct pilot testing to evaluate the infectability of their cell lines and the ability to super-infect without significant toxicity. Also worth mentioning are the informatics analysis: a key challenge in implementing high MOI multiplexing strategies is the need for careful data interpretation. As the number of sgRNAs per cell increases, the likelihood of combinatorial effects and synthetic interactions rises, potentially complicating downstream analysis. While we do not expect issues when conducting large genome-wide screens, guide RNA collisions might be relevant when library sizes are small and can also depend on the expected hit rate. In these cases, future studies should incorporate orthogonal validation methods to ensure robust data interpretation.

In conclusion, this study provides compelling evidence that moderate MOI multiplexing strategies can enhance CRISPRi screen performance, offering a practical approach for researchers seeking to optimize screening efficiency while minimizing resource requirements. By balancing MOI and cell numbers, pooled CRISPR screens can be effectively tailored to address diverse biological questions with improved sensitivity and accuracy. Future research should focus on further refining these approaches, exploring their application to diverse screening paradigms, and integrating complementary technologies to maximize their potential impact.

## Methods

### Cell culture conditions and CRISPRi cell line generation by HDR knock-in into the AAVS1 safe harbor locus

K-562 (ATCC, CCL-243) or HEK 293T/17 (ATCC, CRL-11268) cell are cultured in advanced RPMI (Gibco) with 10%FBS (Gibco) and 1x Pen-Strep (Gibco). Cells are regularly tested for Mycoplasma contamination using MycoStrips (InvivoGen).

To knock in ZIM3-KRAB-dCas9 into the AAVS1 safe harbor locus, K-562 or HEK 293T/17 cells are electroporated with Cas9 ribonucleoprotein (RNP) and a transfer plasmid carrying a ZIM3-KRAB-dCas9 construct flanked by the AAVS1 left and right homology arms. Cas9 RNPs are produced by complexing a human AAVS1-targeting gRNA (targeting sequence: 5’-GGGGCCACTAGGGACAGGAT-3’) to Cas9. The gRNA is ordered as a single guide RNA (sgRNA) from IDT and resuspended at 100 µM in IDTE buffer (pH 7.0). Alt-R HiFi Cas9 enzyme (IDT) and the sgRNA are combined in PBS at a final concentration of 24 µM sgRNA and 20.8 µM Cas9. Complex formation is achieved by incubating the mixture at room temperature for 15 minutes.

2 × 10^6^ HEK 293T or K-562 cells are resuspended in 20 µL of SF Nucleofector solution (Lonza) and to the cell suspension, 5 µL of RNP solution and 1 µg of transfer plasmid DNA was added. The cell suspension is transferred to the electroporation cuvette. Electroporation is done using a Lonza 4D-Nucleofactor X system using the recommended pulse codes for the respective cell lines. Immediately after electroporation, cells are transferred into a 6-well plate with 2.5 mL of pre-warmed medium containing 1 µM HDR Enhancer V2 (IDT). The following day, the culture medium is replaced with fresh medium. 3 days after electroporation, the cells were reseeded in fresh medium containing 20 µg/mL blasticidin (Thermo Fisher Scientific) for selection. 7-10 days post-selection, the cells are sorted for single clones and expanded. The clones are evaluated for functionality and validated.

### Lentivirus production, concentration and transduction

Lentivirus for CRISPR screens and all high MOI transduction experiments is produced in 15 cm dish formats and for validation experiments in 96-well plate formats. 16 × 10^6^ cells per ml or 2 × 10^4^ HEK 293T cells are seeded respectively in 20 mL/150 µL cell culture media with advanced RPMI (Gibco), 10%FBS (Gibco), 1x Pen-Strep (Gibco) and grown at 37 °C, 5% CO_2_. The following day, 10/0.15 µg packaging plasmids and 12/0.25 µg transfer plasmid is mixed with 800/7.5 µl JetPrime buffer and 50/0.7 µl JetPrime reagent (Polyplus) and incubated at room temperature for 10min. The transfection mix is then gently added to the cell culture media. Packaging plasmids are an equimolar mix of psPAX2 (Addgene #12260) and pMD2.G (Addgene #12259). Three days after transfection, cell culture media is collected and cleared by centrifugation (5 min, 500 g, 4 °C). Aliquots from 96-well format preparations are directly frozen at −80 °C before use. Aliquots from 15 cm format are concentrated 50-fold using Lenti-X Concentrator (Takara) according to manufacturer’s recommendation. Concentrated aliquots are resuspended in culture media and frozen at −80 °C before use.

For CRISPR screens and high MOI experiments, cells are transduced using spinfection. Thawed Lentiviral aliquots as well as 8µg/mL polybrene (Milipore) are added to cells in cell culture media. Cells are centrifuged (1000 g, 30 °C, 60 min). Lentivirus and polybrene is left on cells overnight and cells are washed once with cell culture media the following, prior to expanding cells regularly in cell culture media.

### Lentivirus titration using flow cytometry

Lentiviral preps are titrated in 96-well formats. Per well, 5000 cells in 50µl cell culture media (advanced RPMI, 10%FBS, 1x Pen-Strep) and polybrene at a final concentration of 8 µg/mL are mixed with 50µl lentiviral solution. The lentiviral solution is three-fold serially diluted across eleven dilutions. Cell are spinfected as above (1000 g, 30°C, 60min) and the lentiviral solution is replaced with fresh cell culture media the next morning. Cells are cultured for 3 days prior to measuring percentages and mean fluorescent intensities (MFI) of the expressed fluorescent protein. Cells are washed once with EasySep buffer (Stemcell), resuspended in EasySep buffer and measured on an Attune NxT flow cytometer (Thermo Fisher Scientific).

To calculate volumes required to infect at high MOI (>1), the MFI of infected cells in a sample infected at low MOI (∼20% infected cells) is defined as the MFI of a single lentiviral insertion. The MFI is then linearly extrapolated to calculate expected MOI of more concentrated dilutions.

### Copy number quantification using dPCR

To quantify sgRNA expression cassette copy numbers of transduced cells, genomic DNA is extracted with the Monarch Spin gDNA Extraction Kit (New England Biolabs). Purified DNA is normalized to 50 ng/µl. Digital PCR is performed on a QIAcuity One, 5plex device using QIAcuity 26k 8-well Nanoplates and the QIAcuity Probe PCR Kit (Qiagen). Primers and probes are used at final concentrations of 0.8 µM and 0.4µM respectively and 0.25 µl MseI restriction enzyme (New England Biolabs) is added to each 40 µl PCR reaction. RRE targeting primers (5’-AAACTCATTTGCACCACTGC-3’, 5’-AATTTCTCTGTCCCACTCCATC-3’) and probe (5′FAM/-TGTGCCTTGGAATGCTAGTTGGAGT-/3′BHQ_1) and RPL32 targeting primers (5’-CAAGGAAAGACGAGCTGTAGG-3’, 5’-GGGCAGTTGCATCTTCATATTC-3’) and probe (5′HEX/-AGCTGCAGGCAGAAATTCTGGTAGT-/3′BHQ_1) are used to quantify lentiviral insertions and the genomic reference respectively. PCR reactions are run with the following cycling conditions. Initial activation (2 min, 95°C), 2-step cycling with denaturation (15 sec, 95°C) and annealing/elongation (30 sec, 60°C).

### CRISPRi knock-down validation by surface antigen staining

CRISPRi targeting sequences against CD45 (5’-GGTTCTTAGGGTAACAGAGG-3’), CD55 (5’-GAGGCCGGCCCGACGAGCCA-3’), CD71 (5’-GCTCAGAGCGTCGGGATATC-3’), CD81 (5’-GGAGAGCGAGCGCGCAACGG-3’), CD146 (5’-GCAGCAGGCGGCGAGCAAGA-3’) and CD155 (5’-GTCCAGACAAGTGACTGGAG-3’) are cloned into an sgRNA expression vector carrying an eGFP fluorescent reporter and a puromycin resistance gene using Gibson assembly. Lentivirus is prepared and titered individually for each construct. They are mixed at equal titers for combined high MOI transductions. K-562 and HEK 293T cells stably expressing dCas9-Zim3 are transduced, recovered for two days and selected with 2 µg/µl puromycin (Gibco) for three days. After another two days of recovery, cells are stained for flow cytometry in EasySep buffer with anti CD45-Alexa Fluor 700 (Invitrogen), CD58-PerCP-eFluor710 (Invitrogen), CD71-Brilliant Violet 650 (Biolegend), CD81-APC (Biolegends), CD81-APC-Vio 770 (Miltenyi Biotec), CD146-PE (Biolegends) and CD155-APC (Miltenyi Biotec) antibodies. Cells are stained on ice for 30 min, washed twice with EasySep buffer, resuspended in EasySep buffer and measured on an Attune NxT flow cytometer.

### Whole genome CRISPRi sublibrary design and cloning

The design of a whole genome CRISPRi library composed of 24 similarly sized thematic sublibraries is based on a published genome-wide CRISPRi sgRNA libraries^8,31^. Classification relied on the following resources for transcription factors^36^, epigenetic regulators^37^, kinases^38^, phosphatases^39^, cell cycle genes^40^, DNA damage and repair^41^, GPCR^42^, cell surface genes^43^, RNA binding proteins^44^, innate immunity^45^, adaptive immunity^46^, metabolic genes^47^, ion channels^48^, autophagy^49^, apoptosis^50^, ubiquitin^51–56^, cytoskeleton, mitochondria, membrane, ER, golgi, nuclear, extracellular matrix^57^, hydrolase, mRNA processing, ribosomal proteins, proteases, transporter, SLC carriers, cellular structure, olfactory receptors^58^. Genes can be represented in multiple sublibraries in case they fall within multiple classifications. A full list of all sublibrary classifications are found in (**Supplementary Table 3**). Each gene is targeted by three independent sgRNA. In addition to these gene targeting sgRNA, each sublibrary is complemented with 5% non-targeting control sgRNA. The complete annotation of all 24 sublibraries is found in in (**Supplementary Table 3**). Sequences are designed that a single oligonucleotide pool was ordered and amplified with subpool specific primers prior to cloning^59^. Sequences are ordered as a 90K CustomArray oligo pool (Genscript). The 20nt spacer sequences are flanked by 5′-CGATTTCTTGGCTTTATATATCTTGTGGAAAGGACGAAACACC-3’ and 5’-GTTTAAGAGCTATGCTGGAA-3’ sequences containing BsmBI-v2 (NEB) restriction sites and sublibrary specific primer binding sites listed in (**Supplementary Table 3**).

The oligo pool is cloned into a semi-randomly barcoded sgRNA expression vector carrying an mOrange2 fluorescent reporter and a puromycin selection gene. The plasmid is digested with BbsI-HF (New England Biolabs) and purified from a 1% TAE agarose gel using the Monarch DNA Gel Extraction Kit. Each subpool is individually PCR amplified with NEBNext Ultra II Q5 Master Mix (New England Biolabs) and subsequently purified with 2x volume AMPure XP Beads (Beckman Coulter). Purified PCR products are restriction digested with BsmBI-V2 (New England Biolabs) and purified again with 2x volume AMPure XP Beads. A Golden Gate assembly reaction with 20 ng of digested and purified PCR product and 250 ng digested and purified plasmid backbone with 10U Esp3I (New England Biolabs) and 7500U T7 DNA ligase (New England Biolabs) in 50µl reaction were cycled ten times 20°C for 20 min followed by 37°C for 5 min. Each reaction was cleaned up with 1x volume AMPure XP Beads and electroporated into MegaX DH10B T1R (Thermo Fisher Scientific) with a Bio-Rad Pulser II (Bio-Rad) according to the manufacturer’s recommendations. Following electroporation, bacteria were recovered for 1h at 37°C in recovery media and expanded for about 16h at 37°C in LB media with carbenicillin. Plasmid DNA was purified using the NucleoBond Xtra Midi Kit (Macherey-Nagel). To validate the cloning specificity, each subpool library was sequenced on a NovaSeq 6000 (Illumina) at the UCSF Center for Advanced Technologies with read 1 and read 2 lengths of 50 and 50 bases respectively.

### Cell essentiality and drug tolerance CRISPRi screen

Lentivirus of the CRISPRi epigenetic regulator sgRNA sublibrary was produced and titered as described. 700’000 dCas9-Zim3 K-562 cells were spinfected in cell culture media (advanced RPMI, 10% FBS, 1x Pen-Strep) supplemented with 8µg/mL polybrene at anticipated MOI of 0.3,1,2.5,5,10,20 & 30. Cells were expanded for two days and selected with 2 µg/mL puromycin for three days. Infection levels were validated by flow cytometry and digital PCR.

Cells are distributed in duplicates according to the table of conditions below, two days prior to the starting day of the screen. Cell numbers indicated were observed as minimal passaging cell numbers and minimal sampling cell numbers. Each condition was sampled at day 0. Cells were then cultured for 14 days, diluting cells every two days with cell culture media. Imatinib (Sigma) is added at an IC_50_ (230 nM) to the drug treated arm of the screen.

**Supplementary Table 1.** Experimental conditions designed to assess the effects of MOI and cell number on screen performance.

| constant cells |  | constant guide RNA |  | low MOI control |  |
| --- | --- | --- | --- | --- | --- |
| MOI | cells [10E3] | MOI | cells [10E3] | MOI | cells [10E3] |
| 0.3 | 700 | 0.3 | 700 | 0.3 | 700 |
| 1 | 700 | 1 | 700 | - | - |
| 2.5 | 700 | 2.5 | 280 | 0.3 | 280 |
| 5 | 700 | 5 | 140 | 0.3 | 140 |
| 10 | 700 | 10 | 70 | 0.3 | 70 |
| 20 | 700 | 20 | 35 | 0.3 | 35 |
| 30 | 700 | 30 | 23 | 0.3 | 23 |

Cells are collected by centrifugation and frozen at −80°C until further processing. DNA is extracted with the Quick-DNA 96 Kit and quantified on a Nanodrop Spectrophotometer (Thermo Fisher Scientific). All purified DNA is amplified with NEBNext® Ultra™ II Q5 polymerase in 100µl reactions, making sure DNA per reaction does not exceed 20 µg. PCR cycles were 1× (98 °C, 30 sec), 24× (95 °C, 15 s; 65 °C, 20 s; 72 °C, 20 s), 1× (72 °C, 1 min). Forward primers are an equimolar mix of eight staggered primers (5′-AATGATACGGCGACCACCGAGATCTACACTCTTTCCCTACACGACGCTCTTCCGATCT-(N_0-8_)-TTGTGGAAAGGACGAAACACCG-3′), reverse primers include an eight-nucleotide long sample specific index (5’-CAAGCAGAAGACGGCATACGAGAT-(N_8_)-TTGTCCGAGTGACTGGAGTTCAGACGTG-3’) and both are used at a 0.25 µM final concentration. PCR reactions are cleaned up with Mag-Bind; Total Pure NGS beads at a 1.5x concentration and quantified with the Qubit dsDNA HS Assay Kit. DNA is normalized and pooled. Pooled DNA is cleaned up again with Mag-Bind; Total Pure NGS beads (Omega Bio) and diluted to a final concentration of 10 nM. Libraries were sequenced at the CZ Biohub San Francisco on a NovaSeq 6000 (Illumina) with read 1 and read 2 lengths of 70 and 30 bases respectively. Some under-sampled libraries were complemented with a sequencing run on a MiniSeq instrument (Illumina) with the same sequencing conditions.

### FACS-based CRISPRi screen

The CRISPRi screens based on ICAM-1 (CD54) levels was performed in K-562 cell expressing dCas9-Zim3 as follows. A compact genome-wide dual guide RNA expressing CRISPRi library^8,11^ was transduced into 100 × 10^6^ cells at an MOI of 0.3 and 25 × 10^6^ cells at an MOI of 5. Following three days of recovery and four days of puromycin (2 µg/µl) selection, cells were expanded for another four days prior to sorting. Cells are collected by centrifugation (500 g, 5 min, 4 °C), washed once with EasySep buffer, counted and resuspended 10 × 10^6^ cells per ml. Cells are then stained for 30 min on ice with the LIVE/DEAD Fixable Green Dead Cell Stain Kit, for 488 nm excitation (Thermo Fisher Scientific) and CD54-APC antibody (Miltenyi Biotec). Cells are washed twice with EasySep buffer before fixing with the eBioscience Foxp3 / Transcription Factor Staining Buffer Set (Inivitrogen). After fixation, cells are resuspended in EasySep buffer at 20 × 10^6^ cells per ml prior to sorting on a FACS Aria II sorter. Cells are collected by centrifugation after sorting (500 g, 5 min, 4 °C) and cell pellets are frozen at −80 °C prior to further processing.

DNA is extracted as previously described^60^. Cell pellets are resuspended and incubated overnight at 65 °C in 400µl of ChIP lysis buffer (1% SDS, 50 mM Tris, pH 8, 10 mM EDTA) with 16 µl 5 M NaCl. 8µl of RNAse A (10mg/ml - Thermo Fisher Scientific) is added the next day and incubated at 37 °C for 1 h followed by adding 8µl of Proteinase K (20mg/ml - Thermo Fisher Scientific) and a 1 h 55 °C incubation step. Genomic DNA was then isolated with phenol:chloroform:isoamyl alcohol, ethanol precipitated and washed with 70% ethanol. DNA was resuspended in water and quantified on a Nanodrop Spectrophotometer.

Genomic DNA is PCR amplified with NEBNext Ultra™ II Q5 polymerase in five 100µl reactions per sample. PCR cycles were 1× (95 °C, 1 min), 24× (98 °C, 20 s; 63 °C, 45 s; 72 °C, 20 s), 1× (72 °C, 1 min). Forward primers are an equimolar mix of eight staggered primers (5′-AATGATACGGCGACCACCGAGATCTACACTCTTTCCCTACACGACGCTCTTCCGATCT-(N_0-8_)- CCCTTGGAGAACCACCTTGTTG-3′), reverse primers include an eight-nucleotide long sample specific index (5’-CAAGCAGAAGACGGCATACGAGAT-(N_8_)-

GTGACTGGAGTTCAGACGTGTGCTCTTCCGATCTGGCCGCCTAATGGATCCTAG-3’) and both are used at a 0.25 µM final concentration. PCR reactions are cleaned up with Mag-Bind; Total Pure NGS beads at a 1.5x concentration and purified from agarose gel by cutting bands at 750 bp and cleaning them with the Monarch DNA Gel Extraction Kit (New England Biolabs). Cleaned up PCR products are quantified with the Qubit dsDNA HS Assay Kit (Thermo Fisher Scientific). DNA is normalized and pooled 10 nM and sequenced at the Biohub with standard Illumina read 1 (guide RNA 1) and index 1 (sample index) sequencing primers and custom read 2 (guide RNA 2, 5’-GCGGCCAAGTTGATAACGGACTAGCCTTATTTAAACTTGCTATGCTGTTTCCAG-3’) and index 2 (UMI read, 5’-CGTGTGCTCTTCCGATCTGGCCGCCTAATGGATCCTAGTAATCTGCATATCT-3’) sequencing primers. Libraries were sequenced at the CZ Biohub San Francisco on a NovaSeq 6000 (Illumina) with read 1, read 2 and index 2 lengths of 66, 34 and 11 bases respectively.

### Arrayed individual validation experiments

To validate CRISPRi screen hits independently, sgRNA targeting these genes were separately cloned into a sgRNA expression vector by Gibson assembly. A full list of sgRNA sequences is found in (**Supplementary Table 4**).

For drug tolerance validations, test sgRNA were cloned into an expression vector expressing GFP and a non-targeting sgRNA is cloned into a vector expressing mCherry. Cells were lentivirally transduced and selected with puromycin. Cells expressing the GFP-expressing test sgRNA are equally mixed with control mCherry-expressing cells. Ratios of GPF- and mCherry-expressing cells are then measured by flow cytometry under imatinib treated conditions at IC_50_ (230 nM_)_ and compared to untreated controls for twelve days.

To validate genes with effects on ICAM-1 (CD54) levels, test sgRNA were cloned into an expression vector expressing mOrange2. After lentiviral transduction and puromycin selection, live cells were stained with an APC-coupled antibody against CD54 (Miltenyi Biotec) in EasySep buffer for 30 min on ice. After two washes, cells CD54 levels were recorded on an Attune NxT flow cytometer.

### Analysis of CRISPR screens

Read preprocessing and counting of the sequencing data of the sub partitioned CRISPRi library used for the cell essentiality and drug tolerance screen was performed as follows. Forward reads are first trimmed with BBDuk (v. 38.41, parameters: ktrim=l minlen=20 k=10 mink=6 hdist=1 tpe tbo skipr2) to remove the human U6 sequence trailing the spacer sequence (5’-TTGTGGAAAGGACGAAACACCG-3’). Forward and reverse reads are then hard clipped to 20 and 21 bases respectively. Forward reads are counted for perfect matches to the guide RNA library using R (v. 4.5.0) with the packages ShortRead (v. 1.67.0)^61^ and dplyr (v. 1.1.4). To validate sublibrary-specific cloning, guide RNA sequences were counted hierarchically, where reads are first counted only for the sublibrary the guide RNA is expected to originate from to avoid multiple counting of guide RNA represented in multiple of the sublibraries. Barcode and guide RNA diversity as a proxy for unique lentiviral insertion events was calculated using starcode (v. 1.4)^62^. Forward and reverse reads were concatenated prior to running starcode with the --umi-len 21 setting.

Guide RNA level effect sizes were estimated using DEseq2 (v. 1.49.1) comparing untreated day 14 to day 0 samples in each condition. Effect sizes at the gene level are estimated with MAGeCK (v. 0.5.7)^30^ using non-target controls as control sgRNA. Within each condition day 0 samples are compared to untreated and treated samples at day 14 (**Supplementary Table 5-6**).

Data processing of the dual guide RNA CRISPRi library used for the cell sorting screen of ICAM-1 levels was performed as follows. Forward reads are first trimmed with Cutadapt^63^ to remove the mouse U6 sequence trailing the spacer sequence (5’-ATCCCTTGGAGAACCACCTTGTTGG-3’). Forward, reverse and index 2 reads, representing guide RNA 1, guide RNA 2 and the associated barcode, are then hard clipped to 19, 19 and 10 bases respectively. Guide RNA 1 and guide RNA 2 to barcode combinations are individually counted for perfect matches to the guide RNA library and the 41 barcodes using R 4.5.0 with the packages ShortRead (v. 1.67.0) and dplyr (v. 1.1.4). These individual counts are subsequently added yielding for each gene 41 individual barcode counts. These barcode counts are used to estimate gene-level score with MAGeCK again using non-target controls as control sgRNA. Within each condition medium ICAM-1 samples are compared to high and low level ICAM-1 samples (**Supplementary Table 7**).

In our regression, we use DESeq2 estimates because they incorporate standard errors. We run DESeq2 separately on each MOI in the constant cell condition, with the design ∼time. We retain the inferred log2FoldChange as 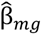 and lfcSE as 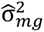 for downstream analysis.

We regress using the formula

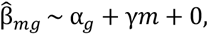

where m indexes the MOI and g indexes the guide. γ is the effect of increasing MOI on the inferred effect sizes. This model is fit using lm in R using weighted least squares, with the weights being 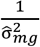.

AUC and ROC-AUC are estimated with the pROC (v. 1.18.5)^64^ package. Accuracy estimated were generated with the caret (v. 7.0-1)^65^ package. Analyses are plotted in R (v. 4.5.0) using the packages ggplot2 (v. 3.5.2) ^66^ & pheatmap (v. 1.0.12) ^67^.

## Acknowledgements

We thank the members of the McManus lab for their feedback and support. We thank Irene Beusch, Diego Calderon, Luke Gilbert and Chris Hsiung for discussion and comments on the manuscript. We appreciate feedback and conversations with Sriram Sankararaman about the analysis. We acknowledge funding support from the Cancer Target Discovery and Development (CTD²) Network (5U01CA272546-03) and the Chan Zuckerberg Biohub. S.O was supported by the Swiss National Science Foundation (SNSF) through an SNF Postdoctoral Fellowship (188001 and 203098). A.X. was supported by the NIH Training Grant in Genomic Analysis and Interpretation T32HG002536. H.P. was supported by the HHMI Hanna H. Gray Fellowship. Some sequencing was performed at the UCSF CAT, supported by UCSF PBBR, RRP IMIA, and NIH 1S10OD028511-01 grants. We thank the Laboratory for Genomics Research, San Francisco for providing the genome wide CRISPRi dual guide RNA library.

## Supplementary Figures

**Supplementary Figure 1:**
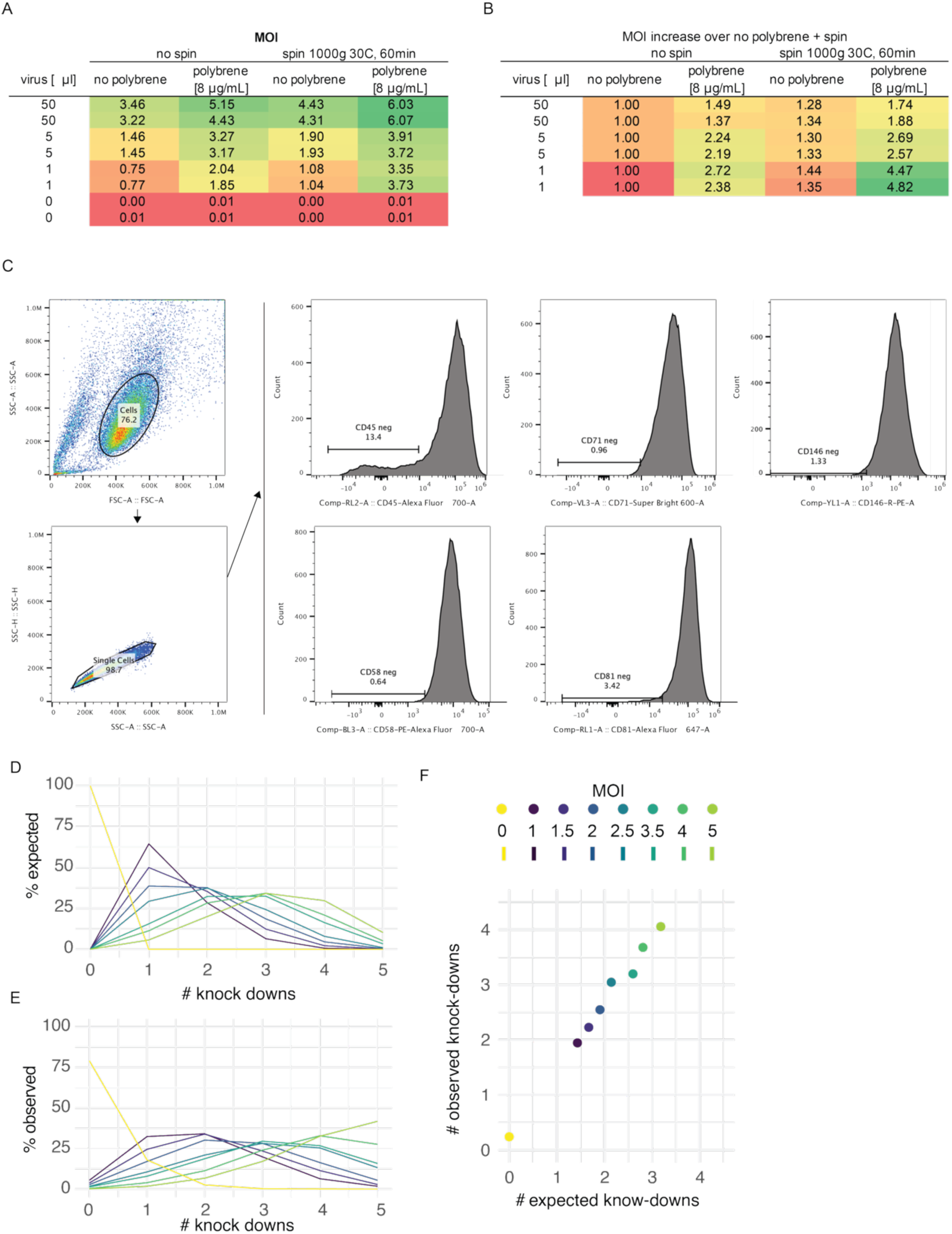
A) Multiplicity of infection (MOI) in relation to variable infection conditions and amounts of virus used for infection of K-562 cells. B) Increase of infection levels relating to the infection conditions in comparison to infecting without the use of polybrene or centrifugation. C) Example of gating strategy to determine percentages knock downs for each of the five observed surface markers. D) Expected numbers of surface marker knock downs relative to the MOI. E) Observed numbers of surface marker knock downs relative to the MOI. F) Average expected and observed numbers of repressed surface markers in respect to infection level upon pooled lentiviral delivery of five sgRNA.

**Supplementary Figure 2:**
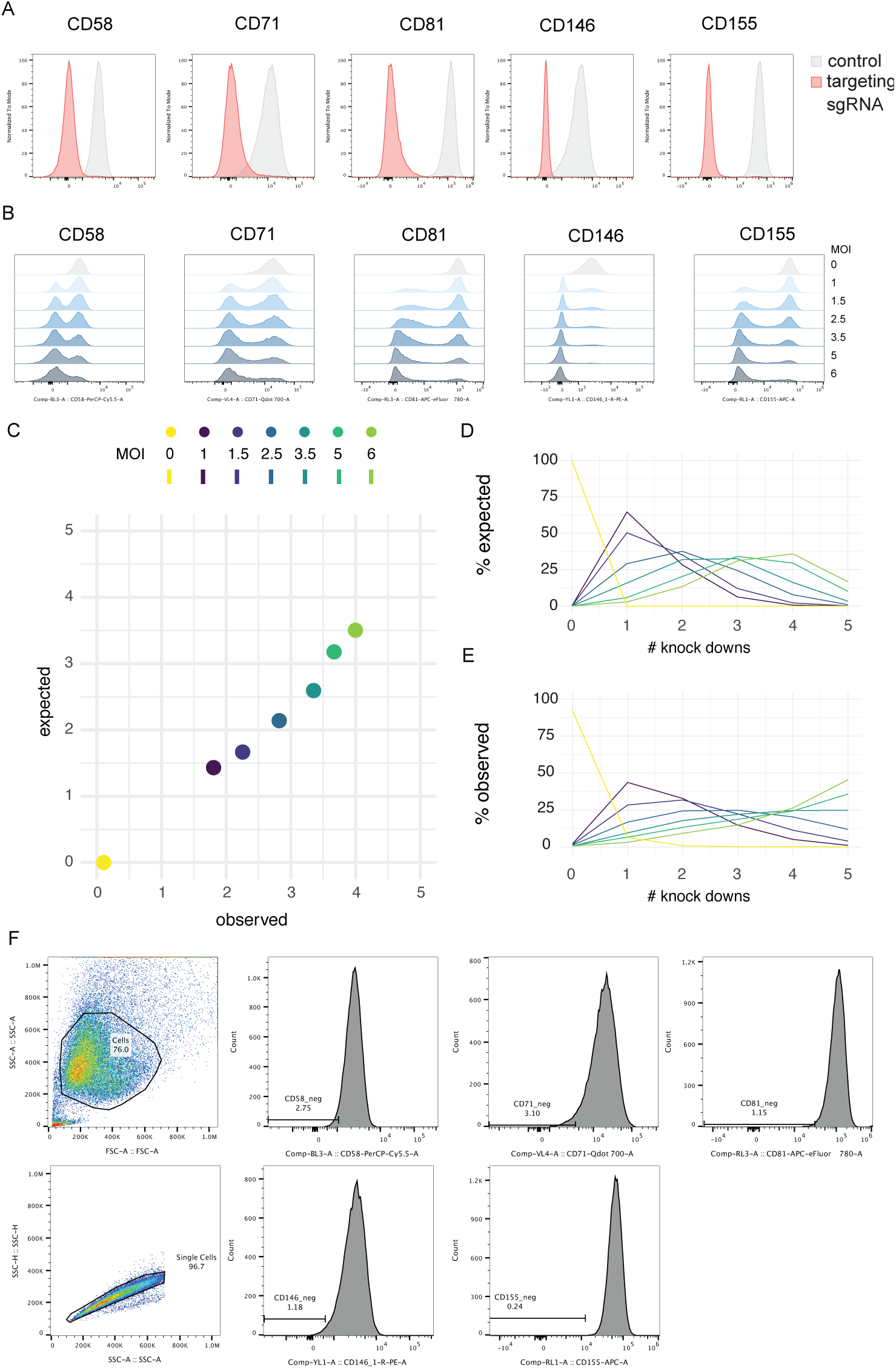
A) Knockdown efficiency of CRISPRi HEK293T cells with individual sgRNA expression against five surface markers. Surface marker expression is measured by flow cytometry of antibody-stained live cells. B) Knockdown efficiency with pooled expression of five sgRNA transduced at increasing multiplicity of infection (MOI). C) Average expected and observed numbers of repressed surface markers in respect to infection level upon pooled lentiviral delivery of five sgRNA. D) Expected numbers of surface marker knock downs relative to the MOI. E) Observed numbers of surface marker knock downs relative to the MOI. F) Example of gating strategy to determine percentages knock downs for each of the five observed surface markers.

**Supplementary Figure 3:**
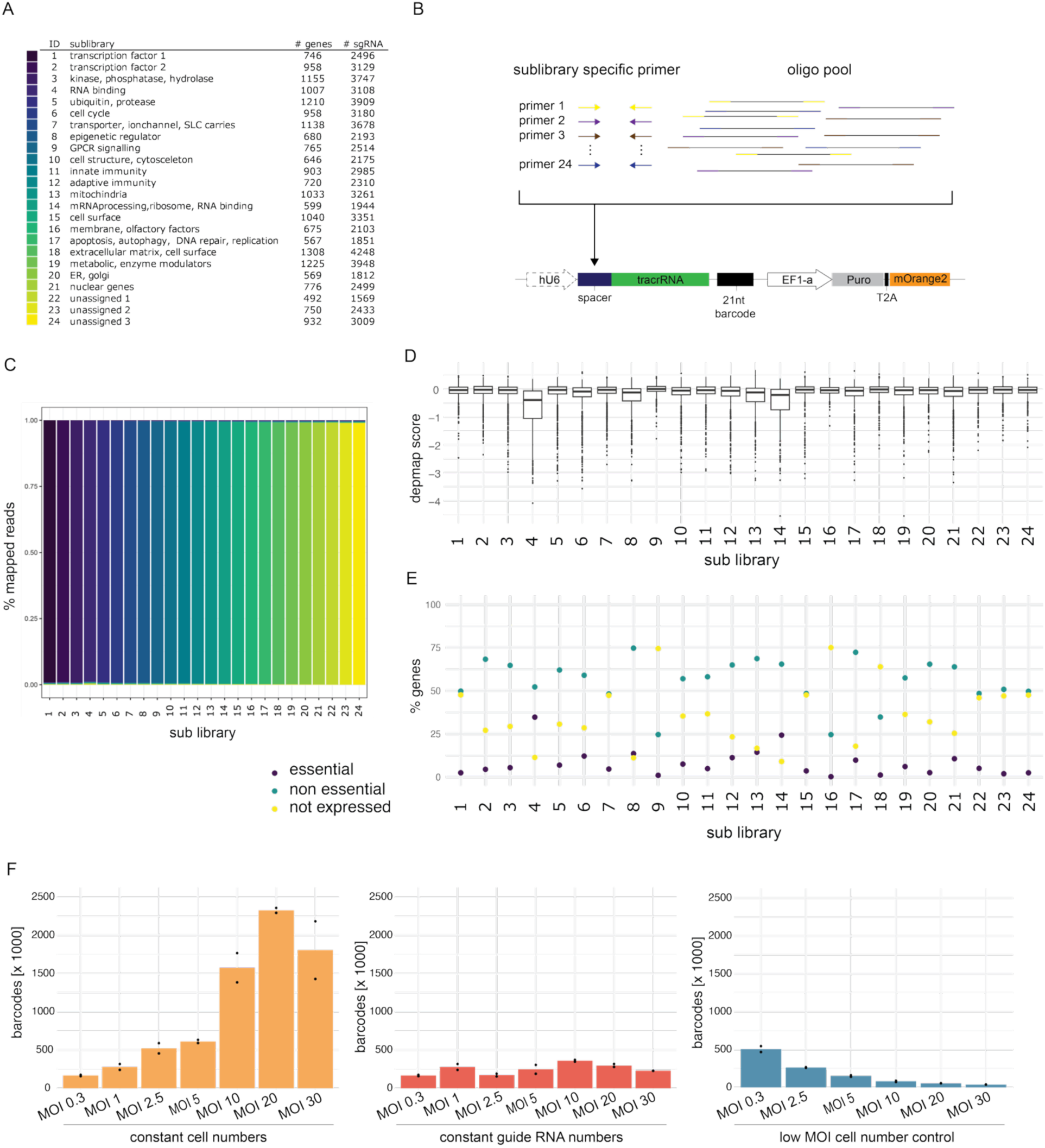
A) Table of gene ontology based CRISPRi sub libraries, numbers of genes targeted and numbers of sgRNA in each sub library. B) Illustration of cloning strategy of CRIPSRi sub libraries using sub pool specific primers on a single oligonucleotide pool. Sub pool specific oligos are specifically amplified with unique primer pairs. C) Specificity of sub pool cloning was confirmed by sequencing the individual pooled library vectors. D) DepMap scores, indicative of cell essentiality in K-562 for genes represented in each sub library. E) Percentage of genes that are cell essential, expressed or not expressed by their representation in the different guide RNA sub libraries. F) Barcode diversity as a measure of the total numbers of lentiviral insertions in each condition. Measured barcode diversity matches anticipated levels and previously established average copy numbers.

**Supplementary Figure 4:**
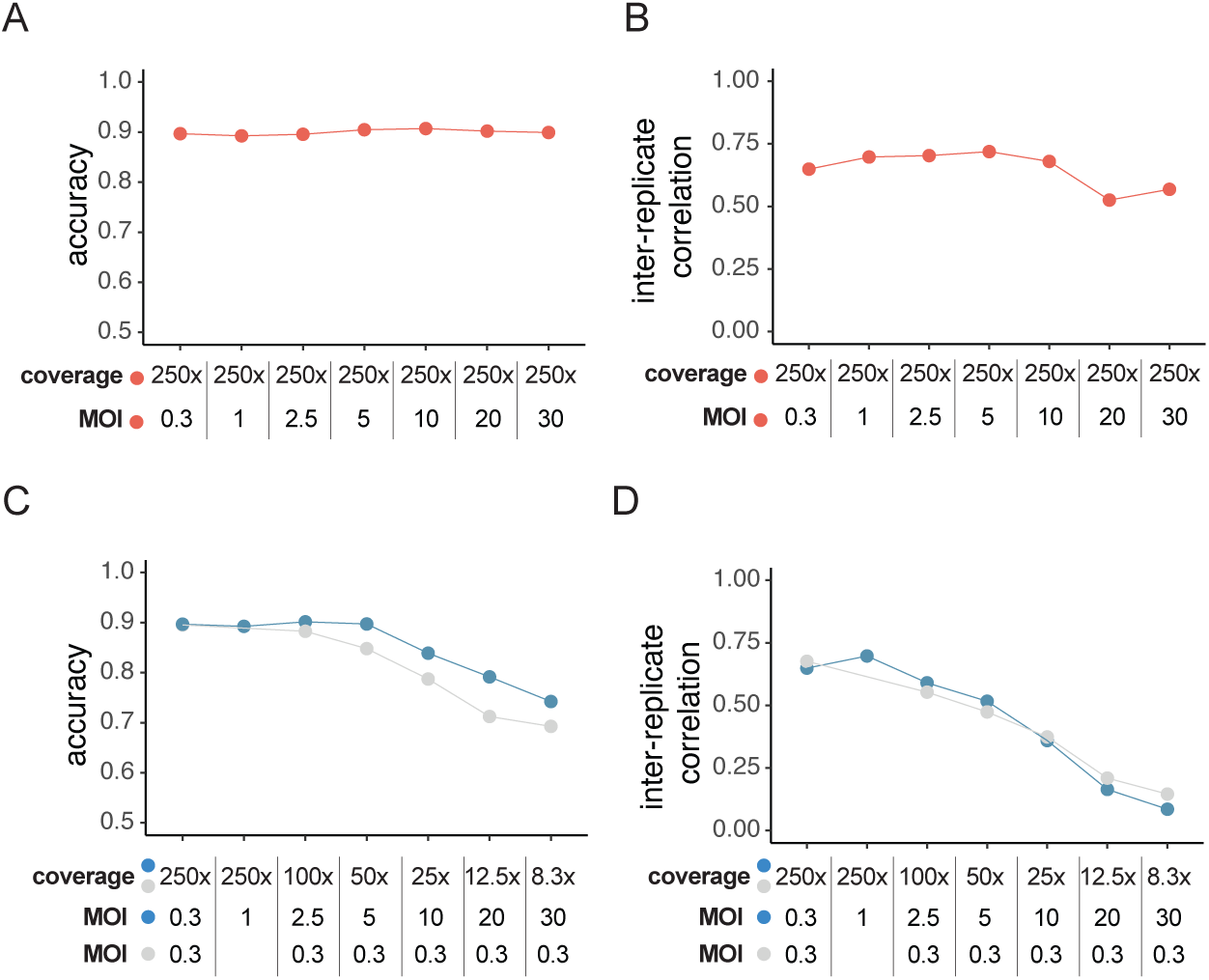
A) Accuracy to classifying cell essentiality effects at the sgRNA level in the constant cells condition (red). B) Inter-replicate Pearson correlation of sgRNA fold changes comparing day 14 and day 0 read counts in the constant cells condition (red). C) Accuracy to classifying cell essentiality effects at the sgRNA level in the constant sgRNA (blue) and low MOI control (grey) conditions. D) Inter-replicate Pearson correlation of sgRNA fold changes comparing day 14 and day 0 read counts in the constant sgRNA (blue) and low MOI control (grey) conditions.

**Supplementary Figure 5:**
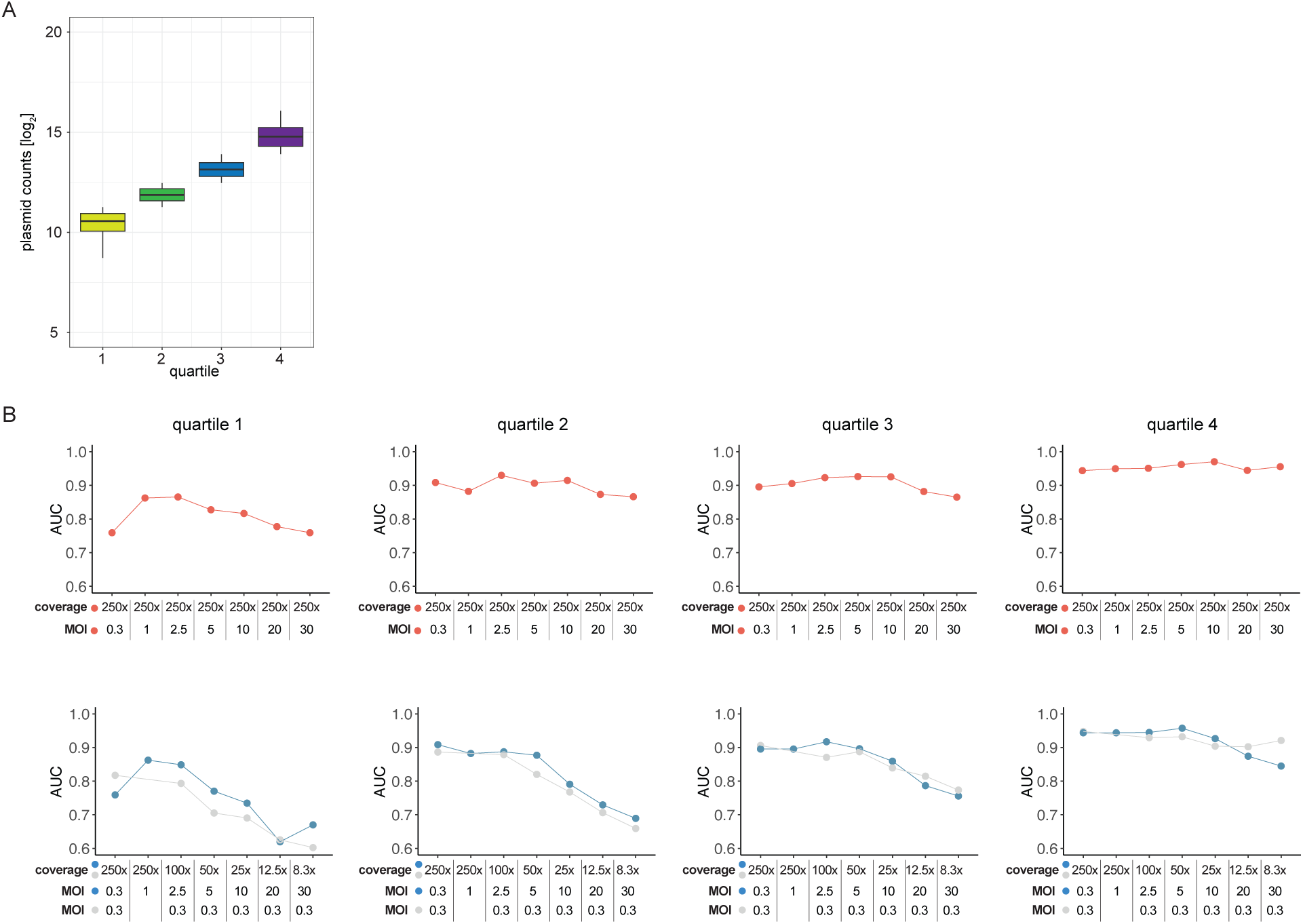
A) Guide RNA are split into four quartiles based on their abundance in the plasmid library. B) Area under the curve (AUC) calculated from ranked sgRNA cumulative distributions. AUC are calculated for sgRNA in the four quantiles split by their abundance in the plasmid library. Constant cells condition (red) and (D) the constant sgRNA (blue) and low MOI control (grey) conditions are plotted separately.

**Supplementary Figure 6:**
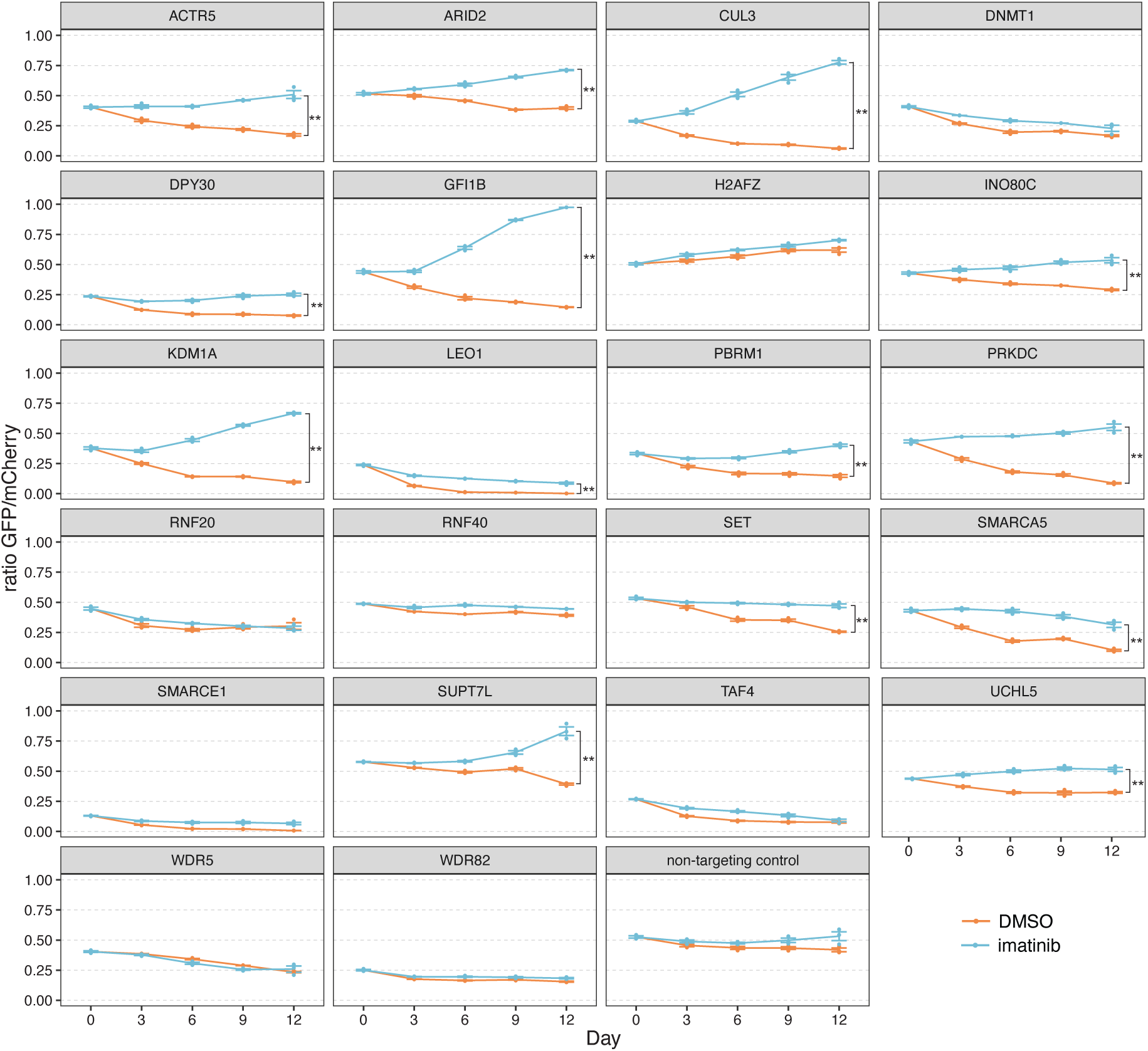
Individual genes are validated for imatinib tolerance. Cells carrying a guide RNA targeting the corresponding genes expressing a GFP marker were mixed with cells carrying a non-targeting guide RNA and an mCherry expressing marking. Paired student t-test comparing the GFP/mCherry ratios at day 12 between imatinib treated (IC_50_, 230nM) and DMSO controls.

**Supplementary Figure 7:**
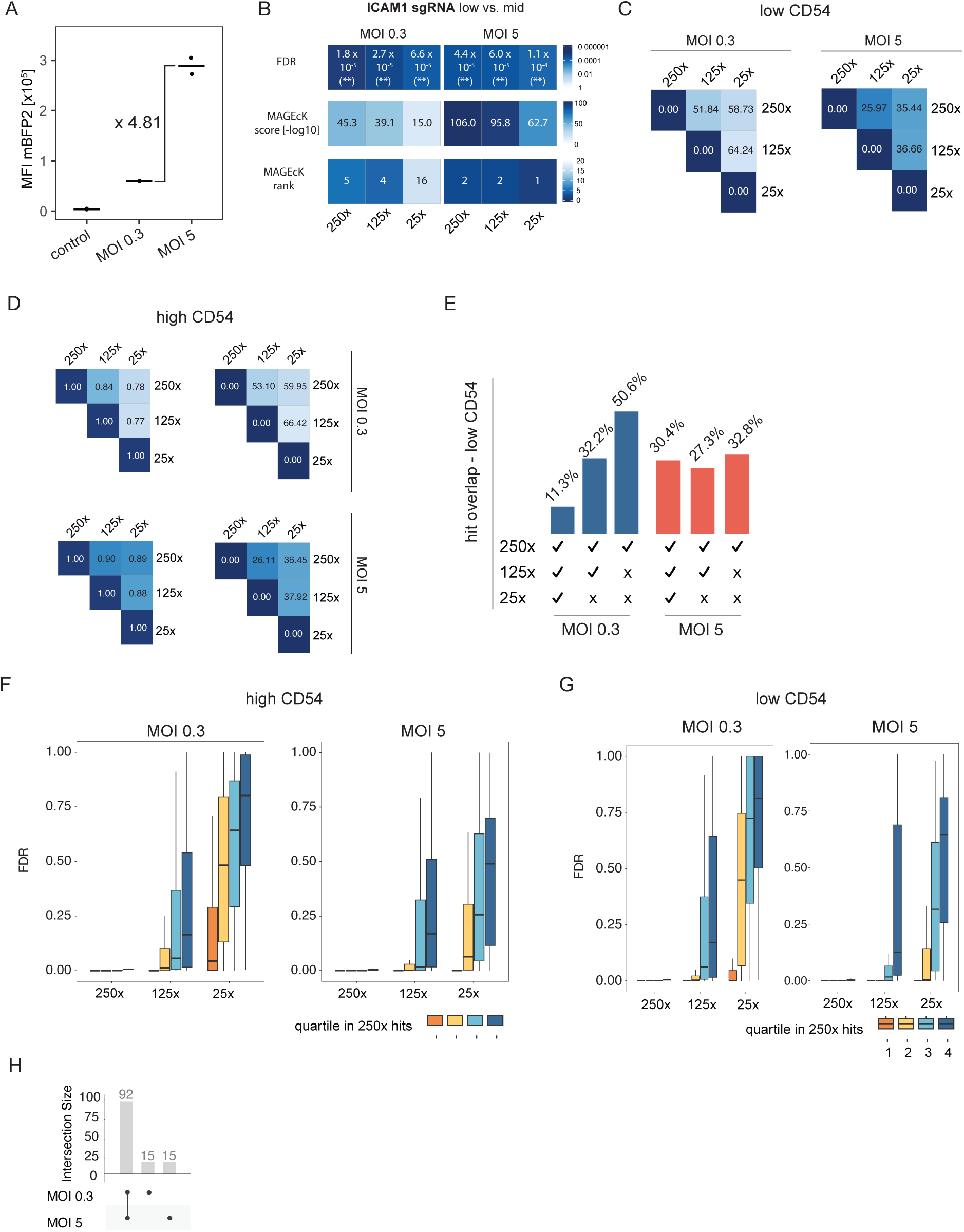
A) Screen results showing guide RNA enrichments based on CD54 levels comparing the bottom 10% of cells to the middle 20% (low vs. mid) or comparing the top 10% to the middle 20% (high vs. mid). False discovery rates (FDR) are shown against log10 transformed MAGeCK scores. Genes are ranked by their MAGeCK score. Results from highest cell number (250X) screens are shown here. Ranking of screen hits based on MAGeCK scores for low vs. mid (B) and high vs. mid (C) comparisons. E) Hits identified in the highest cell number sample (250x) are partitioned into quartiles based on their MAGeCK score in the low CD54 vs. mid CD54 comparison. The most enriched hits are represented in quartile 1 and the least enriched hits in quartile 4. False discovery rates (FDR) of these are shown across different conditions. (Box represents the interquartile range (IQR), spanning from the first to the third quartile, line indicates the median and whiskers extend to 1.5 times the IQR) F) Euclidean distances comparing samples with different cell coverage based on log_2_ fold changes of normalized sgRNA abundances (top). Pearson correlation of the same comparison restricted to genes identified as significant hits in any of the conditions (bottom). G) Percentage of shared hits between cell number settings within the MOI 0.3 and MOI 5 condition in the high CD54 vs. mid CD54 comparison. H) As in (E) but based on the comparison high CD54 vs. mid CD54. Hits identified in the highest cell number sample (250x) are partitioned into quartiles based on their MAGeCK score. The most enriched hits are represented in quartile 1 and the least enriched hits in quartile 4. False discovery rates (FDR) of these are shown across different conditions. True-positive rate (TPR) and false-positive rate (FPR) across all conditions. Sample colors are as in (I).

**Supplementary Figure 8:**
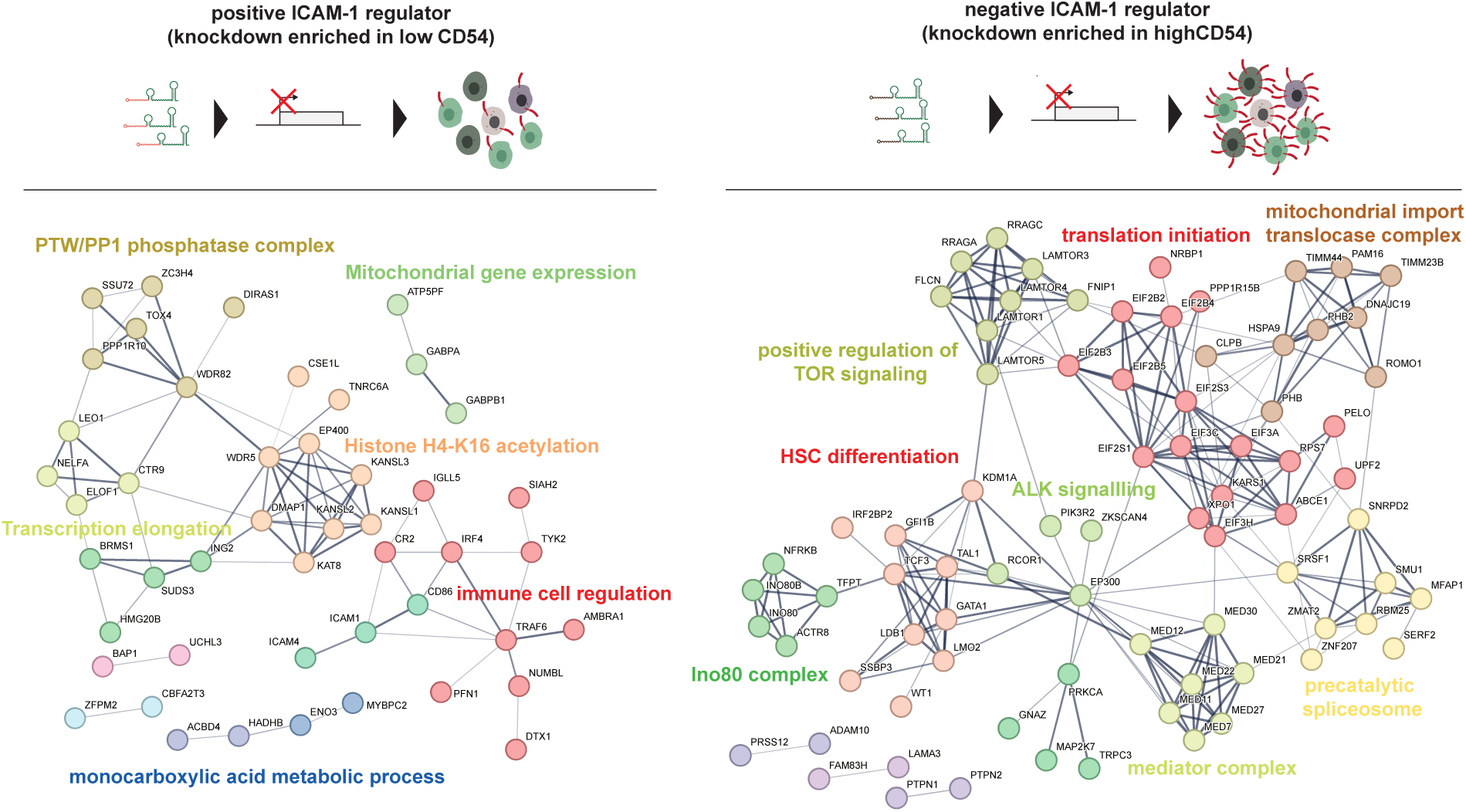
Known gene interactions between genes identified to positively (left) and negatively (right) regulate ICAM-1 levels.

